# Bone Marrow-Derived AXL Tyrosine Kinase Promotes Mitogenic Crosstalk and Cardiac Allograft Vasculopathy

**DOI:** 10.1101/2021.02.04.429773

**Authors:** Kristofor Glinton, Matthew DeBerge, Emily Fisher, Samantha Schroth, Arjun Sinha, Jiao-Jing Wang, J. Andrew Wasserstrom, Mohammed Javeed Ansari, Zheng Jenny Zhang, Matthew Feinstein, Joseph R. Leventhal, Joseph M. Forbess, Jon Lomasney, Xunrong Luo, Edward B. Thorp

## Abstract

Cardiac Allograft Vasculopathy (CAV) is a leading contributor to late transplant rejection. Although implicated, the mechanisms by which bone marrow-derived cells promote CAV remain unclear. Emerging evidence implicates the cell surface receptor tyrosine kinase AXL to be elevated in rejecting human allografts. AXL protein is found on multiple cell types, including bone marrow-derived myeloid cells. The causal role of AXL from this compartment and during transplant is largely unknown. This is important because AXL is a key regulator of myeloid inflammation. Utilizing experimental chimeras deficient in the bone marrow-derived *Axl gene*, we report that *Axl* antagonizes cardiac allograft survival and promotes CAV. Flow cytometric and histologic analyses of *Axl*-deficient transplant recipients revealed reductions in both allograft immune cell accumulation and vascular intimal thickness. Co-culture experiments designed to identify cell-intrinsic functions of *Axl* uncovered complementary cell-proliferative pathways by which *Axl* promotes CAV-associated inflammation. Specifically, *Axl-*deficient myeloid cells were less efficient at increasing the replication of both antigen-specific T cells and vascular smooth muscle cells (VSMCs), the latter a key hallmark of CAV. For the latter, we discovered that *Axl*-was required to amass the VSMC mitogen Platelet-Derived Growth Factor. Taken together, our studies reveal a new role for myeloid *Axl* in the progression of CAV and mitogenic crosstalk. Inhibition of AXL-protein, in combination with current standards of care, is a candidate strategy to prolong cardiac allograft survival.

## INTRODUCTION

Although clinical advances have reduced the incidence of acute cardiac allograft rejection, inflammatory cardiac allograft vasculopathy (CAV) remains a significant contributor to graft failure. CAV is a form of coronary artery disease that is characterized by concentric fibrous intimal hyperplasia along coronary vessels. Evidence of CAV may be found in greater than half of organ recipients during the first 10 years after organ transplant^1^.

In addition to the concentric vascular thickening that characterizes CAV, myeloid cells that include monocytes, macrophages, and dendritic cells are also often found to be elevated^2^. These inflammatory cells have the potential to interact with T cells and also secrete soluble mediators that may contribute to vascular cell activation. In late-stage allografts, intra-graft myeloid phagocytes are associated with worsened clinical outcome^3^.

A key regulator of myeloid inflammation is the TAM (*<u>T</u>yro3/<u>A</u>xl/<u>M</u>ertk*) family of receptor tyrosine kinases. In the case of the protein AXL, this receptor is most notably associated with inflammation resolution. For example, *Axl* gene deficiency, in combination with *Mertk* deficiency, leads to autoimmunity^4^. Although *Axl* may be expressed in many cell types, dendritic cells often exhibit the highest levels^5, 6^. In the setting of transplant, little is known of the role of TAMs. Previous studies by our group have implicated MERTK in tolerance-induction during experimental heart and islet transplantation^7^. However, the causal role of bone marrow-derived *Axl* in cardiac transplantation remains largely unknown. In this context, AXL protein levels have been found to be elevated in multiple instances of solid organ transplant^8^.

Herein we investigated the causal role of recipient and bone marrow-sourced *Axl* during cardiac transplantation. To accomplish this, we performed allogeneic heterotopic heart transplants in *Axl* bone marrow-deficient recipients and monitored key cellular and molecular indices of graft rejection. Our findings newly reveal a detrimental role of *Axl* sourced from the bone marrow, and in cardiac allografts, and offer new insight into how AXL-inhibition may be targeted to reduce myocardial CAV.

## MATERIALS AND METHODS

### Human tissue

This study was approved by the Institutional Review Board at Northwestern University (#STU00012288) and performed in accordance with the Helsinki Doctrine on Human Experimentation. Written consent was obtained from all study participants. Cardiac tissue specimens were obtained from the explanted hearts of adult patients during heart transplantation at Northwestern Memorial Hospital. Explanted hearts were immediately immersed and rinsed with cold cardioplegia solution. Specimens were maintained in cold cardioplegia solution to preserve tissue integrity. Control human samples were taken from normal hearts that were unsuitable for transplant and obtained from patients with severe intracranial hemorrhage. All control samples were obtained from Gift of Hope of Illinois with informed consent. Clinical characteristics of transplant recipients are reported in **Table 1**.

**Table 1.** Cardiac Explant Patient Clinical Characteristics.

|  | **Patient 1** | **Patient 2** |
| --- | --- | --- |
| **Sex** | Female | Male |
| **Weight** | 206 lbs | 256 lbs |
| **Time of First Transplant (Age)** | 4/2004 (17) | 12/2012 (58) |
| **Diagnosis at first transplant** | Non-ischemic cardiomyopathy | Ischemic cardiomyopathy |
| **Time of second transplant (Age)** | 10/29/2018 (31) | 9/16/2019 (64) |
| **Diagnoses at second transplant** | Failing myocardium w/CAV | Failing myocardium w/CAV |
| **Ejection fraction at Second transplant** | 30% | 55% |
| **Comorbidities** | Type II Diabetes, hypertension, hyperlipidemia, obesity | Hypertension, Chronic kidney disease – Concomitant kidney transplant |

### Mice

Eight to twelve-week-old male B6.C-H-2^*bm12*^KhEg (H-2^*bm12*^), BALB/c, and C57BL/6J (B6), mice were obtained from the Jackson Laboratory. ABM mice, described previously^9^, were provided by *M. Javeed Ansari as described*^10^. *Axl-/-* mice were generated as previously described^11^ and littermates of *Axl* heterozygote mating were also utilized in transplant studies. *Cd11c Cre* mice^12^ were crossed to *Axl f/f* mice to achieve >80% mRNA deletion and were from Carla Rothlin at Yale University as previously described^13^. All mice were housed under specific pathogen-free conditions at Northwestern University and approved by the Institutional Animal Care and Use Committee.

### Bone marrow chimeras

Prior to bone marrow transplant, 6-8-week-old mice were subjected to a single lethal dose of irradiation (900 rads). Bone marrow cells were prepared from femurs and tibias harvested from appropriate donors (*Axl+/+* versus *Axl-/-* B6 strains). Femurs were flushed with RPMI medium containing 2% serum. Disaggregated cells were strained through a 40 μm nylon cell strainer and resuspended in serum free RPMI. Irradiated recipients were administered 5×10^6^ cell via tail vein injection. Reconstitution of *Axl-/-* bone marrow was confirmed by flow cytometric analysis of recipient peripheral blood or spleens 4 weeks post-transplant.

### Heterotopic heart transplantation

Abdominal heart transplantation was performed as previously described^14, 15^ and in collaboration with the Northwestern Microsurgery Core. Aged and sex-matched H-2^*bm12*^ or BALB/c mice served as donors to B6 mice. Donor ascending aorta and pulmonary arteries were sutured to recipient abdominal aorta and inferior vena cava to achieve full anastomosis. For complete MHCII mismatch models, mice were administered 2 doses of anti-CD40L (MR1) as specified. For selective inhibition of AXL, R428 (Apex Bio) was reconstituted in hypromellose solution (0.5% hypromellose, 0.1% Tween 80) versus solvent-only control and delivered to mice every other day via oral gavage at 25 mg/kg.

### Echocardiography

Cardiac function was assessed by 2-dimensional M-mode echocardiography using a 25-MHZ probe (Vevo 770; Visualsonics, Toronto, Canada) as previously described^15^. At day 28 after transplant, M-mode images were collected at the level of the papillary muscles, and measurements were made in 3 consecutive cardiac cycles and averaged for analysis. LV-end diastolic and end-systolic dimensions were determined from M-mode tracings and fractional shortening was measured as an indicator of graft function.

### Flow cytometry

Mice were euthanized and grafts were extensively flushed with PBS to remove peripheral cells then excised, minced with fine scissors, and digested with collagenase and DNase at 37°C for 30 minutes as described previously^16^. Graft tissue was subsequently homogenized by pipetting and passed through a 40μm cell strainer. Erythrocytes were lysed and total viable cell numbers were determined by Trypan blue staining. Cells were then incubated with Fc Block (*Biolegend*) for 15 minutes and then labeled with fluorescently-conjugated antibodies for 30 minutes. Flow cytometry was performed on a LSR Fortessa X-20 instrument (*BD Biosciences*) and data were analyzed by FlowJo10.6 software (by *Tree Star*). Macrophages were defined as CD11b^+^Ly6G^-^Ly6C^lo^F4/80^+^. Monocytes were defined as CD11b^+^Ly6G^-^Ly6C^hi^F4/80^lo^. Neutrophils were defined as CD11b^+^ Ly6G^+^. Monocyte-derived dendritic-like cells were CD11b^+^ Ly6G^-^ Ly6C^lo^F4/80^lo^CD11c^hi^MHCII^hi^. T cells were defined as CD45^+^CD3^+^CD19^-^ and further delineated by CD4 and CD8 expression.

### Histology

For whole heart histology, hearts were excised and fixed with 10% phosphate-buffered formalin prior to removal of atria. Fixed hearts were embedded in paraffin and sectioned serially along the lateral axis 6μm apart moving from the base towards the apex. Sections were stained using both a routine hematoxylin/eosin and Verhoeff-Van Gieson’s elastin (*Abcam*) staining protocols.

### Mixed lymphocyte reaction

In a modified mixed leukocyte reaction^17^, we tested the ability of *Axl-/-* dendritic cells to enhance CD4 T cell proliferation. *Axl+/+* versus *Axl-/-* B6 dendritic-like cells were incubated for 4 hours with irradiated total *BM12*^18^ splenocytes and subsequently co-cultured with carboxyfluorescein succinimidyl ester (CFSE)-labeled ABM CD4 T cells, which express TCR specific to BM12^19^. CD4 T cells (non-irradiated) were isolated by magnetic bead selection (*Cell Stemcell Technologies*). Cells were cultured for 72 hours and flow cytometric analysis used to assess proliferation as measured by CFSE dilution.

### Vascular smooth muscle cell (VSMC) culture

Primary vascular smooth muscle cells were harvest from C57BL/6 mice as previously described^20^. Briefly, thoracic aortas were excised, and digested in enzyme solution (collagenase II in Hanks balanced salt solution) for 10 minutes at 37°C. Aortas were then placed in DMEM medium and adventitia carefully removed under a dissecting microscope. Endothelial cells were removed by laterally dissecting the aortas and gently scraping with a damp cotton swab. The remaining tissue was then minced and placed back in enzyme solution and digested for 30 minutes. Disaggregated VSMCs were then washed in DMEM and plated in culture media (DMEM, 20% FBS, 100 u/mL penicillin/streptomycin). Cells were passaged <6 times before use.

### VSMC co-culture and BRDU incorporation

Primary VSMCs were seeded into 48 well tissue culture dishes, allowed to attach overnight, and then serum-starved (0.1% serum; DMEM) for 24 hrs. Synchronized cells were then stimulated with 10% serum, dendritic-like cell (DC)-conditioned media or co-cultured with DCs for 48 hrs. Cells were labeled with APC-BrdU (*BD Biosciences*) for the last 6hrs before harvest. For co-culture experiments, dendritic-like cells were distinguished by elevated CD45 and CD11C staining.

### Statistical analysis

Data represented were mean ± SEM. Significance between groups was calculated by Student two-tailed *t* tests, two-way ANOVA with the Bonferroni post-test or the Logrank (Mantel-Cox) test, as deemed appropriate. A *p* value ≤ 0.05 was defined as significant. All statistical analyses were performed using Prism v8.4.1 (*GraphPad*).

## RESULTS

### AXL is elevated in cardiac allograft vasculopathy

We first set out to determine if AXL protein was present during clinical cardiac transplant rejection. In cardiac allografts from two patients, each diagnosed with varying degrees of cardiac allograft vasculopathy (CAV) (**Table I and Fig. 1A**), we measured increased AXL levels after immunofluorescent staining of cardiac sections and relative to tissue obtained from healthy donor controls (**Fig. 1B**). In parallel, we examined a mouse model of cardiac transplant rejection, in which *B6.C-H-2^bm12^KhEg* (*H-2^bm12^*) hearts were heterotopically transplanted into the abdomens of MHC class II-mismatched *C57BL/6* (*H-2^b^*) mice. In this model, the beta chain of MHC class II molecules differs by a 3 amino acid substitution and is characterized to promote chronic rejection in the absence of immunosuppression. Allografts may develop allo-specific features of CAV, including inflammatory infiltration, progressive concentric intimal thickening, and luminal occlusion following approximately 30 or more days post implantation^19 21^. Consistent with our findings in patients, chronically rejecting mouse cardiac grafts also exhibited evidence of increased AXL staining (**Fig. 1C**).

**Figure 1.**
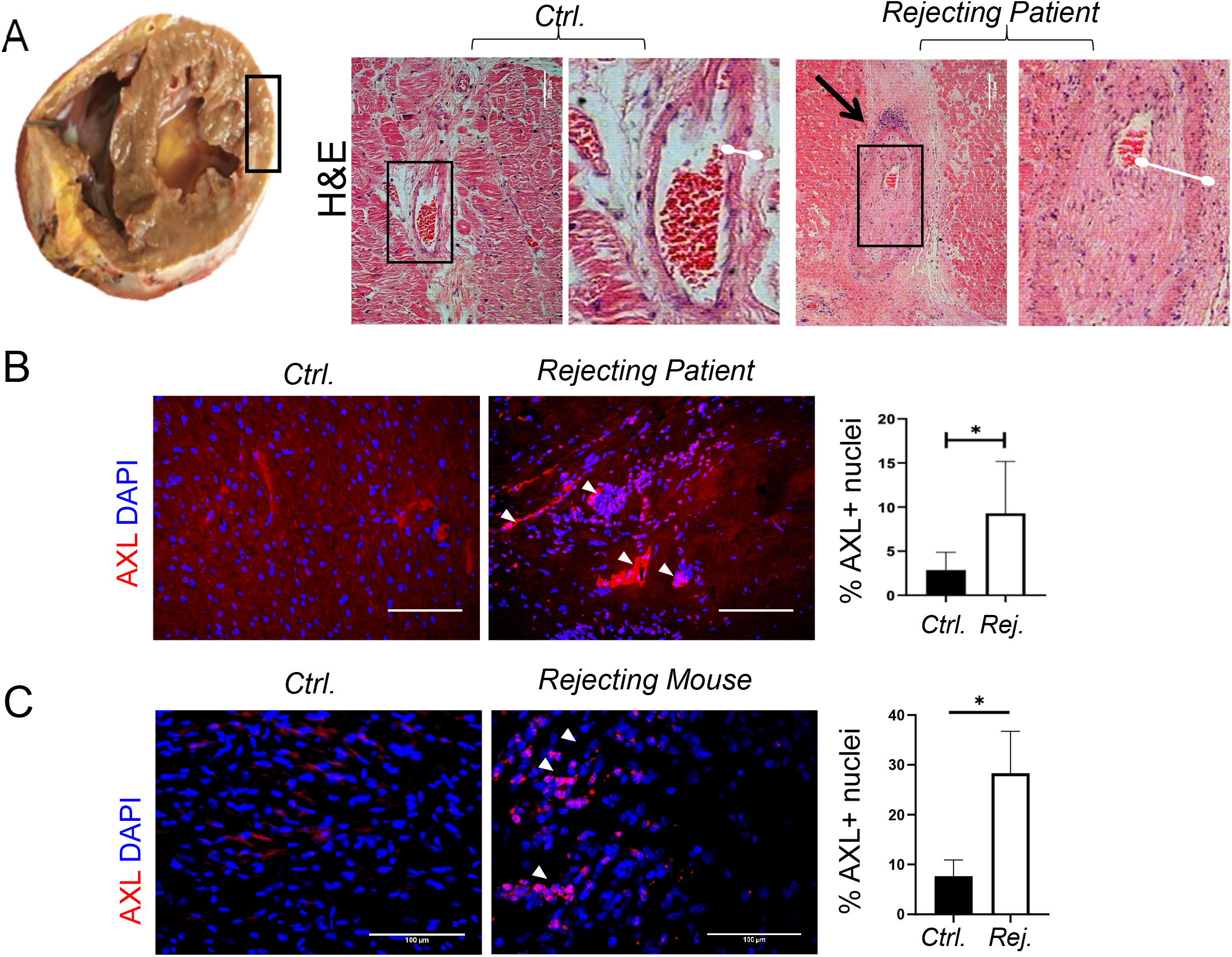
Evidence for elevated AXL protein in rejecting human & mouse cardiac allografts. (**A**) Human heart samples (left ventricle) were procured as described in the Materials and Methods. Rejecting human sample was obtained during the second heart transplant of each patient. Outlined box of specimen indicates analyzed region of interest. Photo-micrographs show control human heart, from healthy donors, but ultimately not transplanted. H&E staining is of explants diagnosed with chronic rejection due to CAV, revealing heightened immune cell infiltration relative to control tissue, indicated by the black arrow. Characteristic evidence of luminal occlusion can also be noted (white bars). (**B**) Immuno-fluorescence staining for AXL staining in rejecting cardiac tissue compared to controls. Enumeration is per regions of interest and quantified as percent fluorescence signal after anti-AXL staining, counter-stained with DAPI for nuclei. White arrowheads indicate positive AXL staining. * indicates p < 0.05 at N=3 vs 3. (**C**) These data are from a BM12→B6 mouse model of CAV. Rejecting grafts (30days) were collected and compared to native mouse hearts from the same recipients as an internal control. Enumeration was calculated as percent signal per nuclei from 4 sections each (Scale bar = 100μm).

### Axl-deficiency improves long term graft survival and function

To ascertain the causal significance of *Axl* to cardiac allografts, we examined transplant rejection in recipients’ deficient for *Axl*. We first transplanted *Balb/c* hearts into *Axl*-deficient *C57BL/6J (B6)* recipients in the presence of costimulation blockade via anti-CD40L/MR1. This approach combines clinically relevant full MHCII-mismatch with co-stimulation blockade to suppress acute graft rejection, yet chronic rejection may occur over time^22^. Improvements in graft survival were documented after transplantation of *Balb/c* hearts into *Axl-/-B6* recipients (mean survival time/MST= 68 days) compared to *Axl+/+* littermates (MST= 60 Days) (**Supplemental Fig. 1**). Importantly, nearly 40% of *Axl-/-* recipients maintained their grafts beyond 100 days, while all *Axl+/+* recipients rejected by day 75. More significant differences were measured after histologic examination as described below.

We next hypothesized a direct role for myeloid *Axl* from bone marrow-derived cells, and because *Axl* expression is not limited to the bone^23^, it was important for us to implement a strategy to focus on this compartment. Therefore, *B6* recipient mice were irradiated and reconstituted with *B6.Axl-/-* versus *B6.Axl+/+* bone marrow at approximately 7 weeks prior to heterotopic heart transplantation (**Fig. 2A**). Importantly, AXL protein was not detectible on the surface lymphoid T or B cells before or after bone marrow transplant of *Axl+/+* cells, in contrast to *Axl-/-* cells. (**Supplemental Fig. 2**). To study the direct consequences of bone marrow-derived *Axl*, in the absence of immune co-stimulation blockade, we utilized the aforementioned *BM12* model of chronic allograft rejection^19^. Longitudinal assessment of grafts by manual palpations revealed that recipient bone marrow-derived *Axl*-deficiency resulted in significantly improved clinical scoring (**Fig. 2B**). We further assessed indices of graft function, including graft ventricular fractional shortening (FS), using echocardiography at 4 weeks post-transplantation. In agreement with manual assessments, grafts in *Axl*-deficient recipients maintained significantly higher FS at this time point (**Fig. 2D, E**). Graft mean survival time (MST) was also >20% improved in *Axl-/-* recipients relative to *Axl+/+* animals (**Fig. 2C**). Importantly, and consistent with the therapeutic relevance of our findings, selective AXL-inhibition showed significant efficacy at improving graft viability after complete MHCII-mismatch (**Fig. 3**).

**Figure 2.**
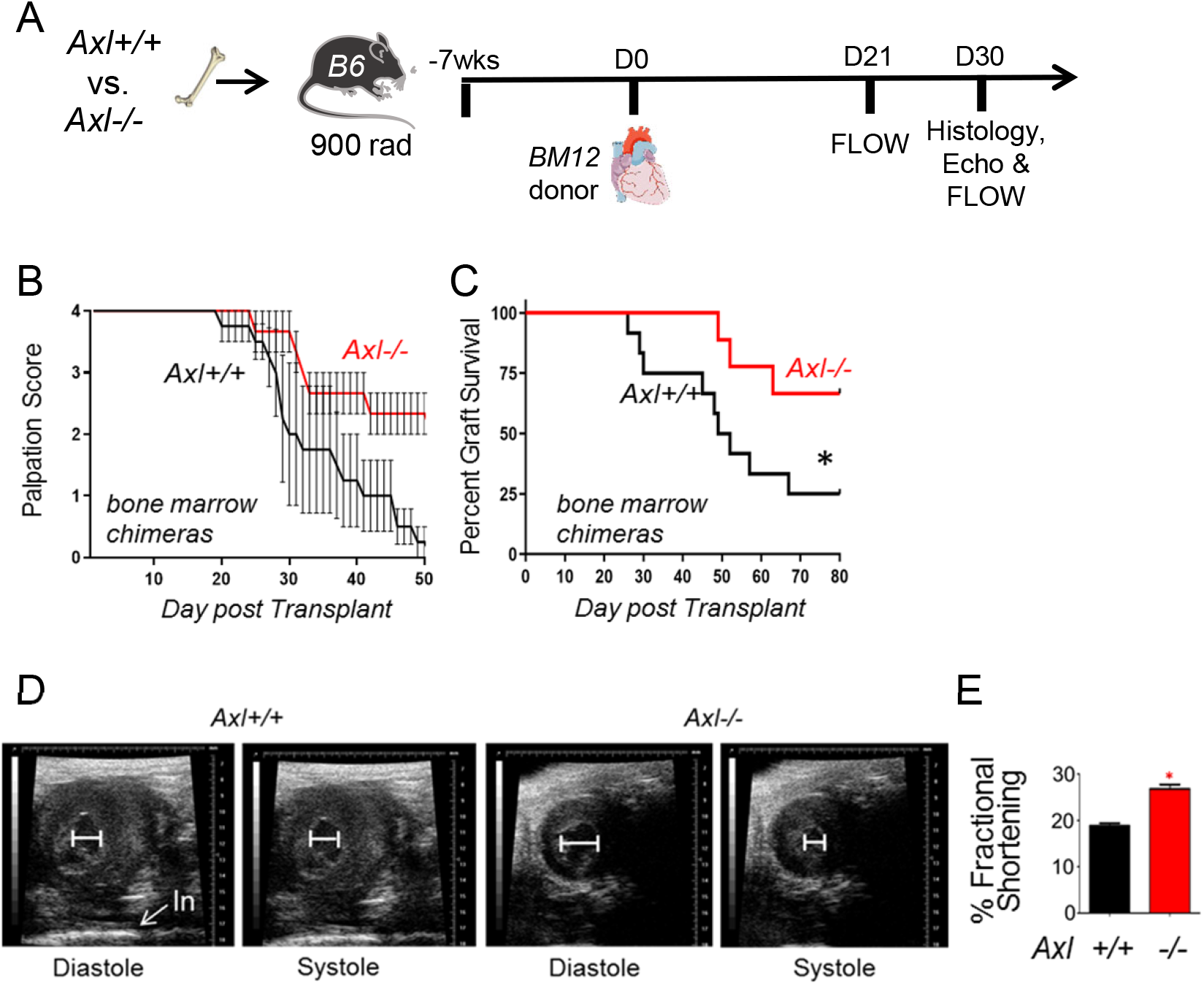
Bone marrow-derived *Axl* promotes chronic allograft rejection. (**A**) Irradiation and bone marrow transplant were carried out by 7 weeks prior to heart transplantation in C57BL/6 (B6) mice. *BM12* hearts were implanted into B6 recipients and graft palpation was utilized to quantify clinical score after palpation (**B**) and percent graft survival (**C**) from indicated genotypes. In addition to manual palpations, contraction of graft ventricles was measured by echocardiogram (**D**) at 28 days post-transplant (**E**). N = 12 versus 10 * P<0.05. “In” indicates intestinal landmark after heterotopic cardiac implantation.

**Figure 3.**
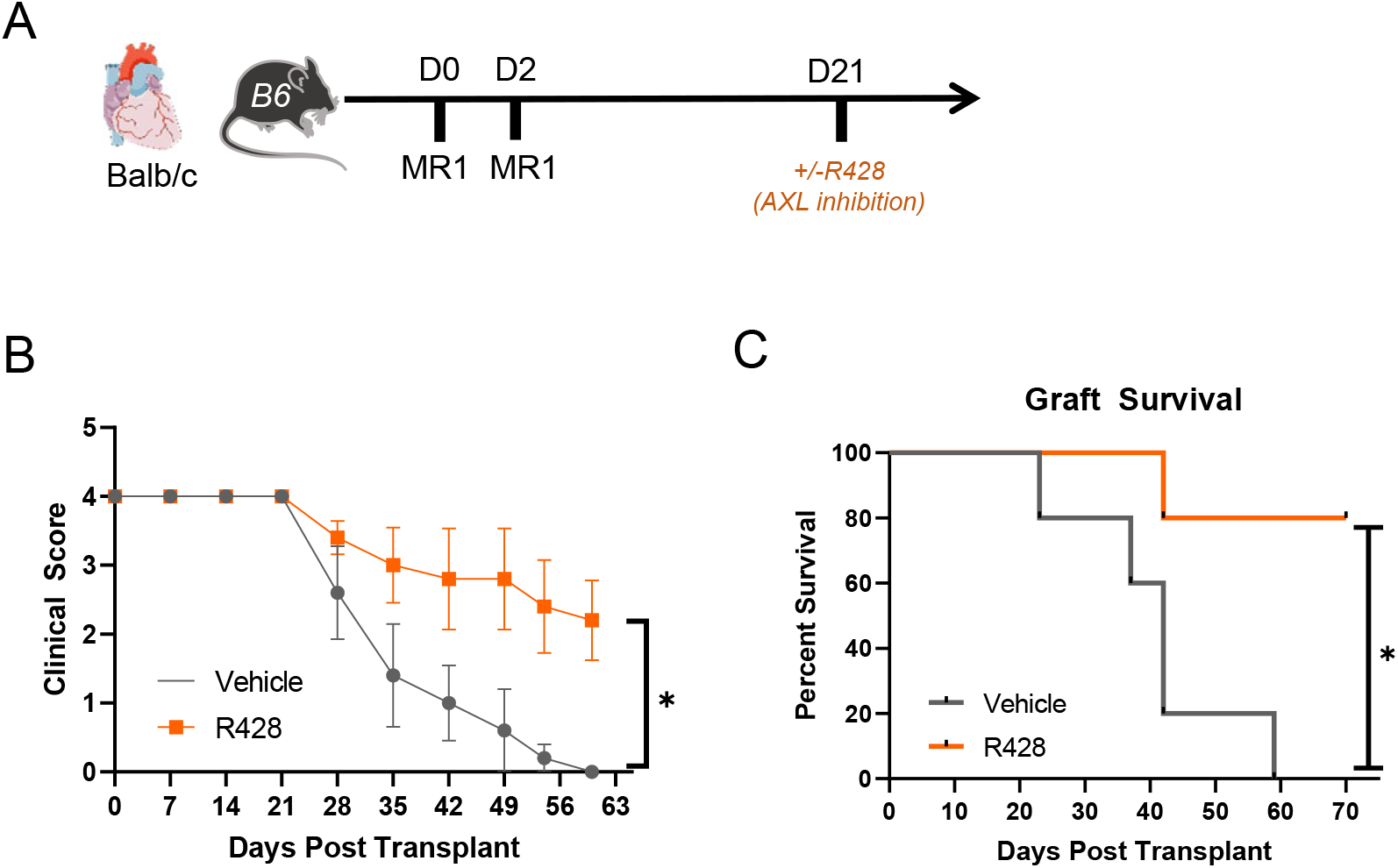
AXL-inhibition improves allograft clinical score and survival. (**A**) B6 recipients received anti-CD40L (MR1) on D0 and D2 post-transplant. The AXL-inhibitor R428 or vehicle was administered every other day via oral gavage to B6 recipients 3 weeks after transplant. (**B**) Depicted are clinical scores after experimental *Balb/c to B6* heart transplantation. * p < 0.05 2way ANOVA (n=5 per group) (**C**) Overall graft survival was also measured after infusion of the R428 AXL inhibitor. * p < 0.02 Mantel Cox Test.

### Reduced CAV and diminished allograft inflammation in the absence of Axl

A significant cause of long-term allograft rejection is inflammatory CAV. Vasculopathy in rejecting hearts was indicated increased luminal occlusion in *Axl+/+* animals relative to *Axl-/-* recipients (**Fig. 4**). Reduced intimal thickening was also measured after transplantation of full MHCII-mismatched *Balb/c* hearts into *B6* recipients (**Supplemental Fig. 1B**). Furthermore, allografts from *Axl-*deficient recipients also exhibited an overall trend of reduced numbers of innate immune cells, including Ly6G^+^ neutrophils, Ly6c^HI^ monocytes, F4/80^+^ macrophages, and CD11b^+^ dendritic-like cells (**Fig. 5, Supplemental Fig. 3**). Lymphocyte CD19^+^ B cells and CD4 and CD8 T cells were also reduced (**Fig 5, Supplemental Fig. 4**). Similar reduction in lymphocyte infiltrate was observed in recipients with targeted deletion of *Axl* in CD11c+ cells (**Supplemental Fig. 5**). The only immune population to trend upward in our panel of *Axl*-deficient recipients were tolerogenic Foxp3^+^ T regulatory cells^24^ (**Fig. 5 and Supplemental Fig. 6**). Interestingly analysis of splenic immune cell content revealed heightened numbers of neutrophils and monocytes in *Axl*-deficient recipients, suggesting reduced mobilization to inflamed tissue (**Supplemental Fig. 7**).

**Figure 4.**
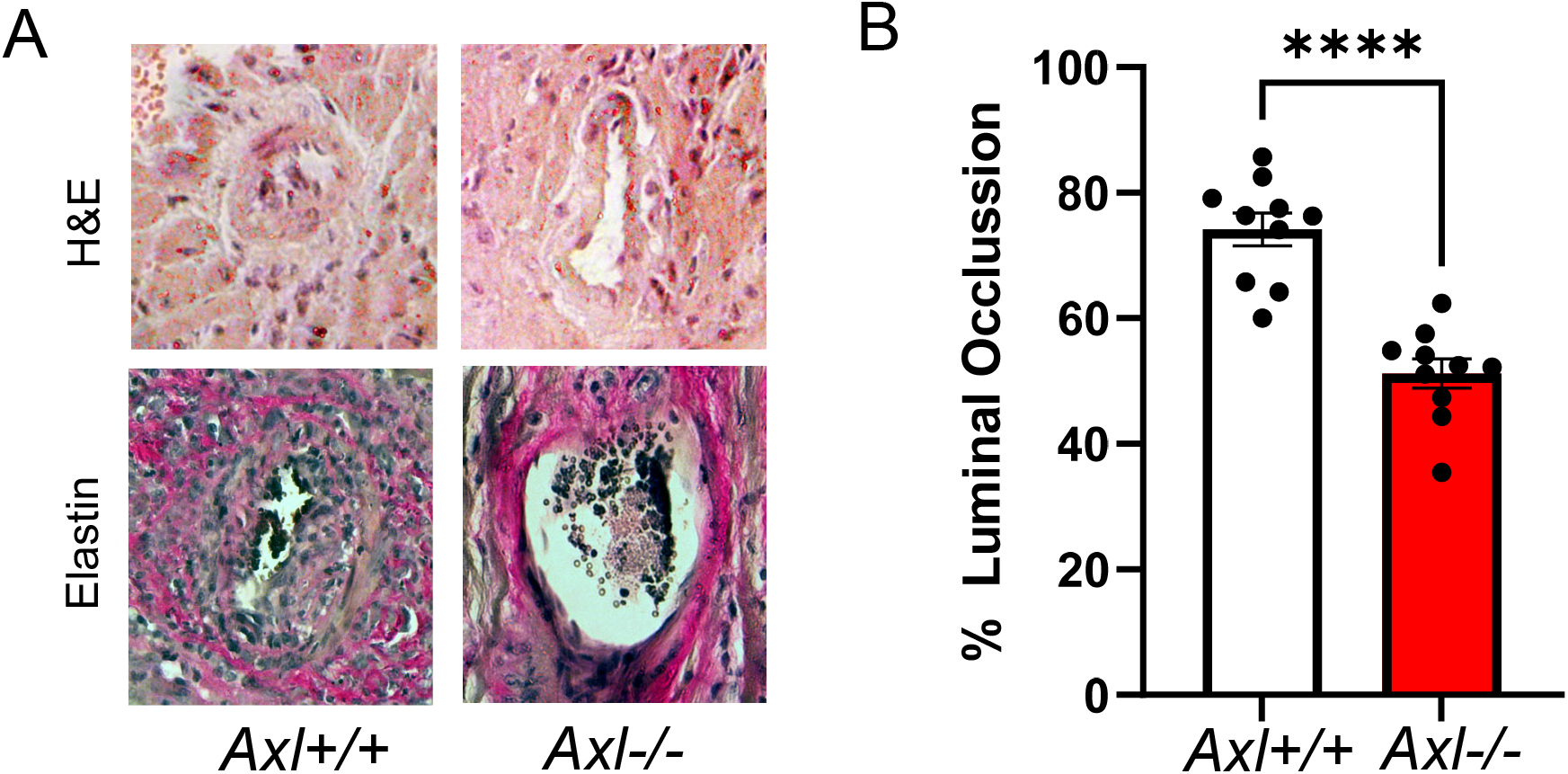
Bone marrow-derived *Axl* deficiency reduces markers of cardiac allograft vasculopathy. (**A**) Histologic analysis of allografts after *BM12→B6* heterotopic heart transplant. (**B**) Quantification of percent luminal occlusion, calculated as (IEL-LA)/IEL × 100. lEL= Internal elastic lamina area, LA = Luminal area. N=10 vs 10. ****p < 0.01.

**Figure 5.**
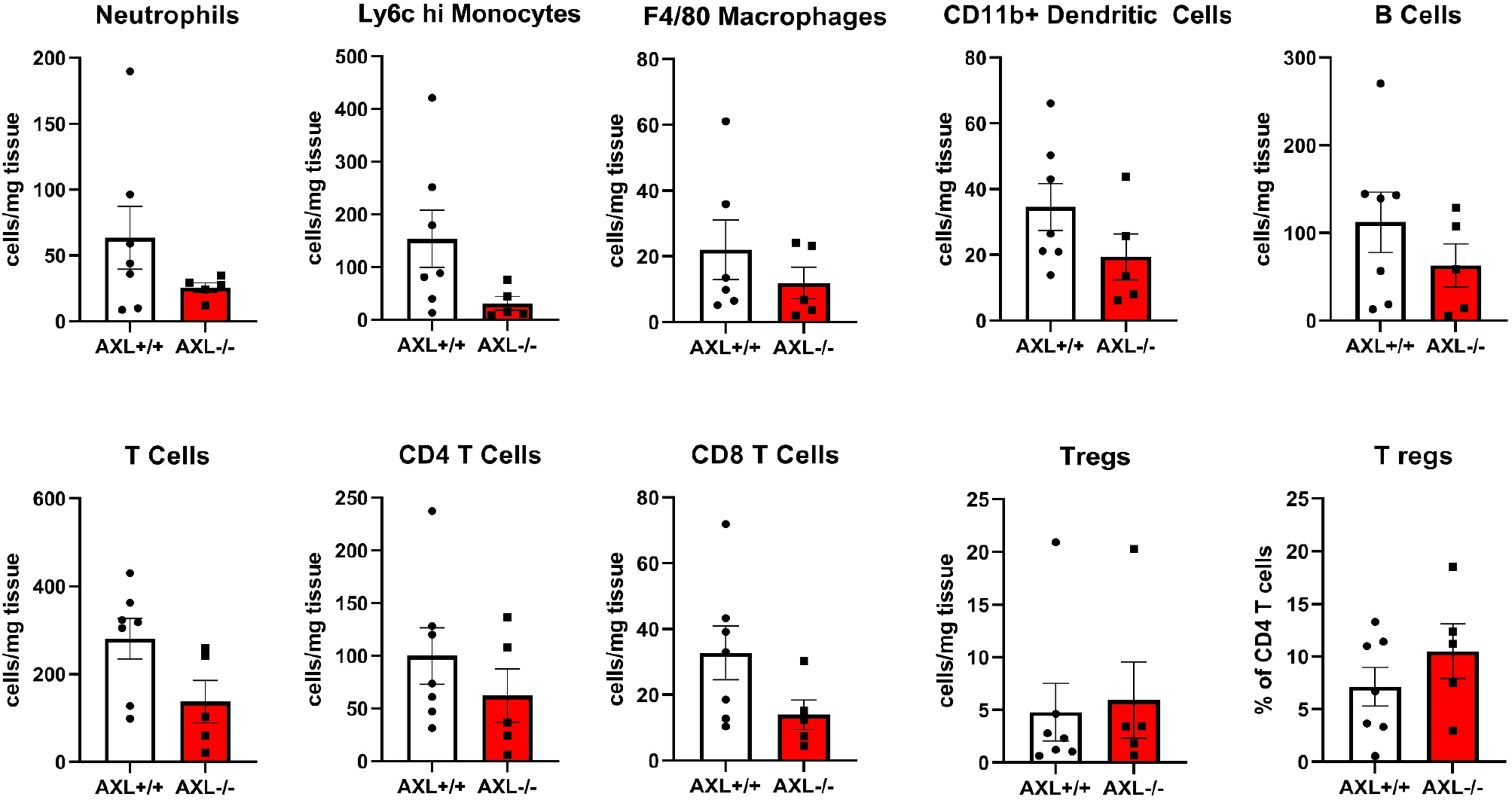
Reduced allograft inflammatory cell accumulation in the absence of bone marrow-derived *Axl*. BM12 cardiac allografts were extracted and subjected to flow cytometric analysis from B6 *Axl+/+* versus *Axl-/-* recipients. Neutrophils were Ly6G^+^ Macrophages were F4/80^HI^. For T cell subsets, CD45^+^CD3^+^ cells were gated to distinguish CD4 and CD8 levels. T regs were further identified as CD4^+^CD25^+^Foxp3^+^ Data represents the combination of 3 independent experiments where N=5-7 mice per group. *p<0.05for *Axl*+/+ versus *Axl*-/-.

AXL is known to be highly expressed on dendritic cells^25^, where it has been found to directly correlate with MHCII expression^13^. Indeed, AXL co-localized with HLA in human sections (**Fig. 6A**), and MHCII in mice, 21-30 days post-transplant (**Fig. 6B**). We also performed flow cytometric analysis of allografts, and discovered that graft CD11c^+^ cells of *Axl+/+* animals displayed significantly higher levels of both MHCII and costimulatory molecule CD86 relative to *Axl-/-* recipients (**Fig. 6C**). Additionally, cultured *Axl-/-* dendritic-like cells exhibited reduced MHCII and CD86 expression by flow cytometry (**Fig. 6D**), as markers of myeloid cell activation. In a similar fashion, grafts transplanted into animals with CD11c specific deletion of *Axl* also displayed reduced MHCII expression in this subset. Interestingly these cells also displayed increased levels of inhibitory ligand PD-L1, which has also been linked to enhanced Treg development^26^ (**Supplemental Fig. 8**).

**Figure 6.**
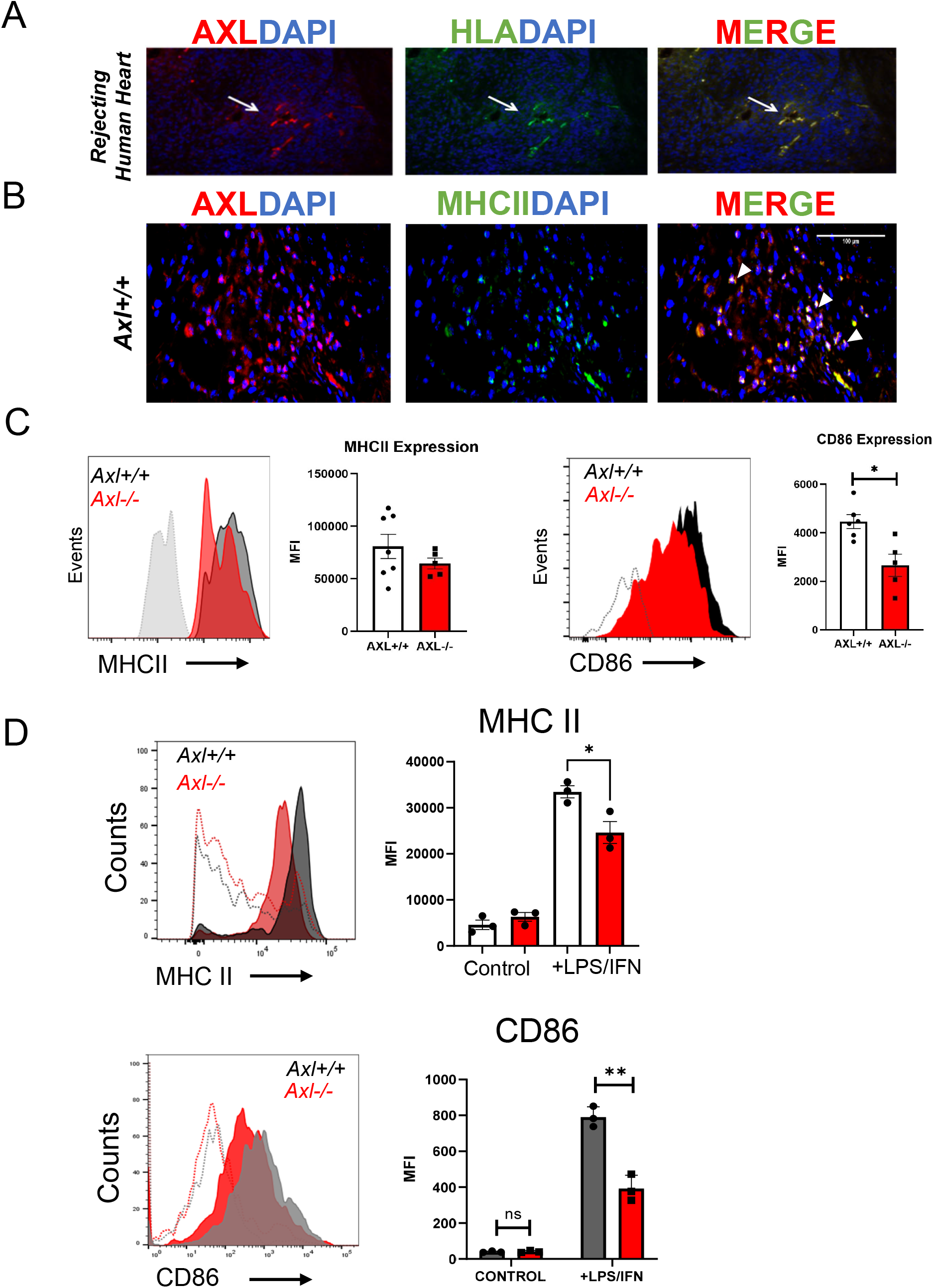
Evidence that elevated AXL coincides with markers of cellular activation. (**A**) Immunofluorescence with anti-AXL and counter-stained with DAPI for nuclei from patients. (**B**) In mice: Immunofluorescence for anti-MHCII followed by image merge of AXL and MHCII signal. (**C**) Cardiac allografts were analyzed by flow cytometry for levels of dendritic cell surface MHCII and costimulatory molecule CD86. Dotted line indicates FMO flow cytometry control for each condition. Data represents the combination of 2-3 independent experiments where N=10 mice per group. (**D**)In culture, *Axl+/+* versus *Axl-/-* myeloid cells were generated and stimulated with LPS and IFN then surface levels of MHCII and costimulatory molecule CD86 measured by flow cytometry. Histograms and MFI are representative of 2 independent experiments where N=3 mice per group. *p < 0.05 *Axl+/+* versus *Axl*-/-.

### Myeloid Axl-deficiency leads to suppressed cell proliferative crosstalk

We were next interested in determining cell-intrinsic mechanisms by which recipient *Axl* in bone marrow-derived cells may promote transplant rejection and vasculopathy, Given the link between *Axl* and cancer progression^27^, we first considered that *Axl* could be involved in cell proliferative pathways involved in advancing cardiac allograft rejection, such as T cell mediated activation^28^ and vascular smooth muscle cell proliferation^2^.

In the case of T cell activation after transplantation of *H2-Ab1^bm12^* hearts into *C57BL/6* recipients, donor bm12 MHC alloantigen^18^ is recognized via the direct antigen presentation pathway^9^. Therefore, we hypothesized a mechanism by which recipient myeloid *Axl* promotes paracrine signals during allograft inflammation to enhance T cell activation. To model this scenario, we cultivated a modified mixed leukocyte reaction whereby *B6.Axl*-deficient myeloid cells (relative to *Axl+/+* control) were incubated with irradiated BM12 splenocytes and BM12-specific ABM T cells. Anti-bm12 ABM T cells directly recognize MHC BM12 alloantigen^19^. From this culture, we subsequently measured antigen-specific ABM CD4 T cell proliferation. As hypothesized, cultures exposed to *Axl-deficient* dendritic-like cells exhibited a significantly reduced ability to stimulate ABM CD4 T cells (**Supplemental Fig. 9**).

In parallel, a signature feature of CAV is concentric intimal hyperplasia, driven primarily by migration and proliferation of vascular smooth muscle cells (VSMCs). Chronic low-level inflammation has been postulated to promote VSMC phenotype switching and proliferation, and intra-graft myeloid cells are associated with worsened clinical outcome^3^, however the source of such agonists has not been completely investigated. To determine if *Axl* might also regulate VSMC proliferation, we generated co-cultures in which primary cell-cycle synchronized VSMCs from *Axl+/+* animals were cultured in the presence of *Axl+/+* versus *Axl-/-* dendritic-like cells (**Fig. 7A-C**). We found that in the presence of *Axl+/+* myeloid cells, a significant percentage of VSMCs incorporated the proliferation marker BrdU and were found to be in the S phase of mitosis. In contrast, co-culture with *Axl-/-* myeloid cells exhibited significantly blunted replication and BrdU incorporation. This *Axl*-dependent difference also manifested when VSMCs were incubated in the presence of conditioned dendritic-like cell media (**Fig. 7D-F**), consistent with the mechanism being non-cell contact dependent.

**Figure 7.**
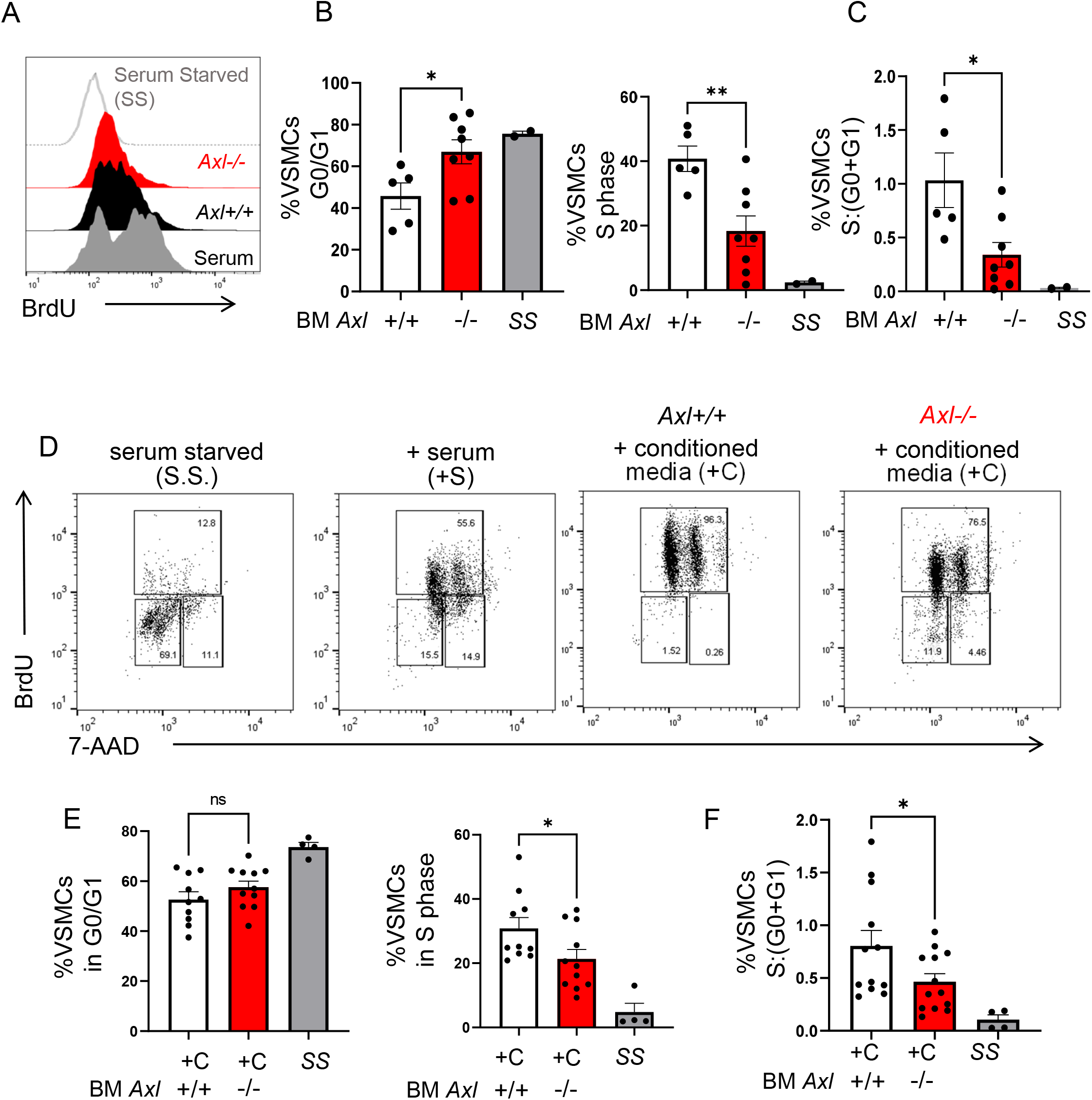
Reduced vascular smooth muscle cell (VSMC) proliferation when cocultivated with *Axl*-deficient myeloid cells. Primary murine B6 *Axl+/+* VSMCs were cocultivated with bone marrow-derived *Axl+/+* versus *Axl-/-* myeloid cells. (**A**) VSMCs were cultivated with nucleoside analog bromodeoxyuridine (BrdU) for flow cytometric analysis as an indication of cell proliferation. (**B**) To measure VSMC cell cycle, 7-amino-actinomycin D (7-AAD) was also added to quantify double-strand DNA. As depicted, myeloid *Axl*-deficiency was associated with elevations of VSMCs that were arrested in the G0 and G1 phase of interphase and elevated in S phase. (**C**) Ratio of VSMCs in S phase relative to VSMCs in G0 and G1 phase. Data represents the combination of 3 independent experiments where N=5-8 mice per group. *p<0.05 *Axl-/-* versus *Axl*+/+. (**D**) Replication of VSMCs was synchronized by culture in serum free medium and subsequently cultured for 4 days with serum compared to conditioned media from bone marrow-derived *Axl+/+* versus *Axl-/-* dendritic cells. Markers of VSMC proliferation and quiescence were measured by flow cytometry after addition BrdU and 7-AAD was also added to quantify double-strand DNA to assess cell cycle stage. (**E**) Quantification of flow cytometric data revealed that media from *Axl*-deficient myeloid cells led to ~15% reduction in VSMC proliferation as determined by reduced BrdU positive cells in the S (synthesis) phase of interphase, concomitant with a higher percentage of cells arrested in the G0/G1 phase. (**F**) Ratio of VSMCs in S phase relative to VSMCs in G0 and G1 phase. Data represents the combination of 3 independent experiments where N=10 mice per group. *p <0.05 *Axl+/+* versus *Axl*-/-.

In further considering the contribution of *Axl* crosstalk with VSMCs, we discovered evidence for myeloid cell expression proximal to VSMCs in our histologic analysis of rejecting allografts as evidence of possible myeloid-VSMC crosstalk (**Supplemental Fig. 10**). We then asked if VSMC proliferation in co-culture with *Axl+/+* myeloid cells required the VSMC mitogen PDGF^29^. Indeed, we found that blocking-antibodies specific for PDGF significantly impaired VSMC BrDU levels when in culture with Axl+/+ conditioned myeloid cell media, newly characterizing a myeloid cell PDGF-dependent VSMC proliferation pathway. However, in parallel this phenotype was not recapitulated with media sourced from *Axl-/-* cells (**Fig. 8A**). Indeed, *Axl* was required for accumulation of PDGF in conditioned media (**Fig. 8B**). Thus, *Axl* is required for PDGF-dependent VSMC proliferation. Taken together, our findings reveal complementary evidence, both cell-intrinsic and *in vivo*, for *Axl*-dependent promotion of CAV-related pathways after cardiac transplantation.

**Figure 8.**
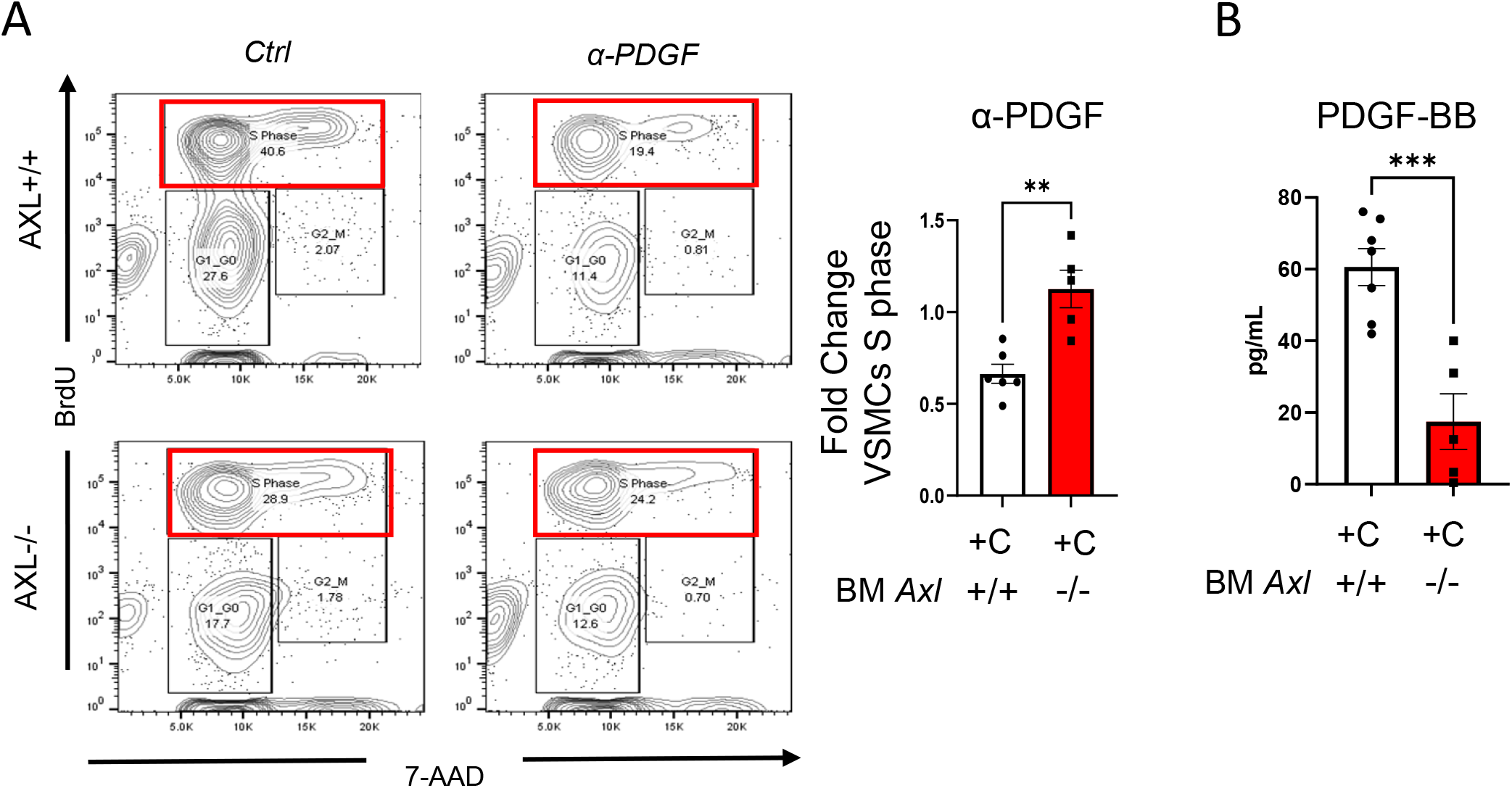
Axl is required for PDGF-dependent VSMC proliferation. Bone marrow-derived myeloid cells were cultured in GM-CSF and IL-4 for 7 days. Non-adherent cells were collected for further culture. VSMCs were plated and following serum starvation to synchronize the cell cycle, cells were cultured with conditioned media from myeloid cells ±PDGF blocking antibody (20μg/mL). For cellular proliferation analyses, BrdU was added 24 hours prior to harvest. Cells were analyzed by flow cytometry (**A**) and S phase fold-change (after blocking antibody treatment) of VSMCs quantified. (**B**) *Axl* is required for PDGF accumulation in cell supernatant. PDGF-BB isoform in conditioned media of GM-CSF/IL4 derived myeloid cell was quantified by ELISA. Data represents the combination of 2 independent experiments in which N=5-7 mice per group. ** p= 0.002 ***p=0.001.

## DISCUSSION

Our findings newly implicate a role for *Axl* tyrosine kinase, specifically sourced from a bone marrow-derived compartment, in the promotion of cardiac allograft vasculopathy. At the cellular level, our results single out a common mitogenic pathway by which *Axl* may enhance transplant rejection mechanisms. In particular, vascular smooth muscle cell/VSMC activation is a hallmark of transplant vasculopathy, and our data indicate that *Axl* promotes VSMC proliferation. Interestingly, the implicated mechanism by which this occurs is through *Axl-*dependent production of Platelet-Derived Growth Factor. A working model of our findings is depicted in **Supplemental Fig. 11**.

Separately, the results of our experiments also implicate *Axl* in enhanced alloantigen-reactive T cell proliferation. In the case of the BM12 model employed in this study, this may be explained by donor allograft passenger immune cell infiltration into lymph nodes, followed by direct recipient Tcell allo-recognition and paracrine *Axl*-dependent pro-proliferative crosstalk. In this context, it will be interesting to test the contribution of *Axl* on donor passenger immune cells. Another non-mutually exclusive scenario is in the setting of complete MHC mismatch. For example, *Axl*-dependent increase in MHC II and co-stimulatory ligand expression may stimulate T cell activation through indirect allo-antigen presentation. Taken together our studies are consistent with potential complementary pathways by which *Axl* may regulate allograft survival.

As aforementioned, multiple studies have highlighted the anti-inflammatory and protective role of TAM family members, including *Axl*. Together, TAMs act as pleiotropic inhibitors of the innate immune response and prevent autoimmunity^4^. Thus, our findings for a pro-inflammatory role of *Axl* in the transplant setting, relative to many studies, may appear to some as a surprise. However, emerging evidence supports the concept that individual TAM members act through divergent signaling pathways^6^. For example, and relative to the TAM MERTK, AXL protein uniquely interacts with ligands Gas6 (which is expressed in allografts)^30^ and Protein S to afford differing thresholds of activation, as monitored by tyrosine kinase phosphorylation^31^.

*Axl* may be expressed on non-myeloid cells, such as endothelial cells and smooth muscle cells^32–34^. However, our strategy specifically targeted *Axl* sourced from the bone marrow and did not find evidence of AXL protein on lymphoid cells. Nevertheless, we cannot discount potential non-myeloid effects of pharmacological interventions that utilize non cell-specific AXL-inhibitors. Separately and during after MHC mismatch, myeloid bone marrow-derived dendritic cells capture and transport antigen to lymph nodes where they interact with T cells and T regulatory cells^35, 36^, including during organ transplant^35, 37^. Post-transplant, donor or “passenger” antigen-presenting cells are eventually replaced by those of the recipient and may interact directly with inflamed graft cells^38^. The underlying mechanisms by which recipient dendritic cells contribute to chronic rejection are not fully elucidated. In bone marrow-derived dendritic-like cells, *Axl*-dependent phagocytosis has been shown to enhance antigen cross-presentation and T cell activation to apoptotic cell-associated antigens^17^. In addition, *Axl* has been associated with altered MHC levels^13, 33^, as we now newly document in the setting of transplant, although the underlying mechanisms remain unclear and are a focus of future studies.

A key indicator of CAV is concentric intimal wall thickening of the vasculature, resulting in vascular occlusion and eventually graft failure. Wall thickening involves active migration and proliferation of vascular smooth muscle cells (VSCMs). VSMCs have been shown to undergo phenotypic switching, triggered by inflammatory mediators^39^. Our data are consistent with the possibility that VSMC activation during transplant rejection is regulated by AXL in subendothelial leukocytes, which may be found proximal to activated VSMCs. Future studies are required to elucidate how AXL may specifically regulate PDGF levels, or other AXL-dependent growth factors.

Our study, like other murine models of transplant^40^, has limitations to consider in its clinical relevance. This includes the nature of the *BM12* model, which may not reflect transplants with greater MHC variance. Nevertheless, this model was useful in our studies to understand inflammation that emerges during allo-recognition, as well as mechanisms by which AXL directly regulates transplant rejection and in the absence of potentially confounding immunosuppression. In parallel, we utilized a complete MHC mismatch transplant model, administering costimulatory blockade (MR1 anti-CD40 Ligand) to *C57BL/6* recipients whom had received *BALB/c* transplants. In this scenario, characteristics of VSMC hyperplasia were similarly reduced in *Axl-/-* mice. In addition, alternative pathways of chronic rejection that are mediated by NK and B cells, that latter which contributes to chronic antibody-mediated rejection^41^, were not addressed in these studies. For example, it has been reported that AXL protein signaling is necessary for optimal human NK cell development^42^. Considering evidence that NK cells may contribute to CAV in an IL-6 dependent manner^43^, we cannot exclude contributions of this pathway to our findings. Furthermore, *de novo* donor-specific antibody production often correlates highly with the development of CAV^44^. This predominantly involves activation and expansion of B cells. Interestingly, unchecked AXL phosphorylation on B cells has been reported in progression of lymphoblastic leukemias^45^. It remains to be seen if *Axl* has a direct effect on B cell maturation in solid organ transplant, however one may speculate that inefficient T cell priming could impede B cell responsiveness. It is also interesting to note that a separate report described GAS6-mediated AXL signaling in T regulatory cells to enhance associated FOXP3 and CTLA4 expression^46^. In our study, we did note a trend of increased graft infiltrating T-regulatory cells.

Taken together, our studies reveal a novel role for AXL in response to allograft transplant. Future efforts are necessary to elucidate mechanisms of AXL-dependent crosstalk with T cells, as well as the identity of potential other paracrine factors that may regulate dendritic-like cell or macrophage cross talk with VSMCs. It also remains to be seen if these mechanisms are organ-specific or may be further applied to other solid organ transplants. Our current observations suggest that the therapeutic targeting of AXL in chronic rejection may be warranted to reduce vasculopathy and enhance longevity of grafts.

## Acknowledgements

This work was supported by the National Institutes R01HL139812, R01HL122309, and an AHA postdoctoral fellowship to 18POST33960228 KG. We appreciate the assistance of Lisa Wilsbacher, M.D. Ph.D., FAHA, Anna Huskin, RN, BSN, CCRC, Program Development Manager in the Clinical Trials Unit of the Bluhm Cardiovascular Institute and Patrick McCarthy, MD, the Executive Director of the Bluhm Cardiovascular Institute.

## Disclosure and Conflict of Interest Statement

The authors have nothing to disclose. The authors do not have relevant financial activities or intellectual property outside the submitted work.

---

**Supplemental Figure 1.**
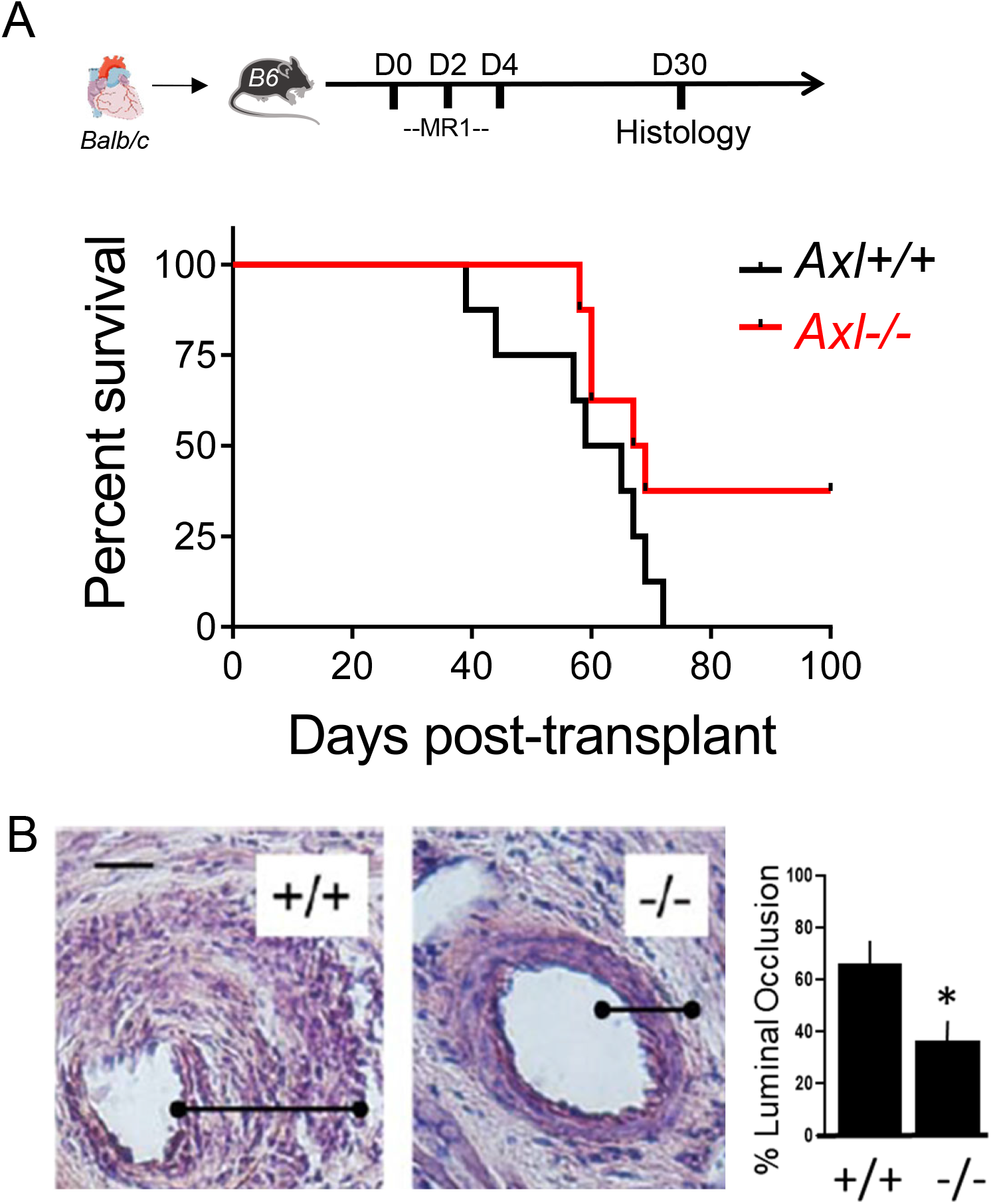
Full MHC mismatch model of transplant in the absence of *Axl*. (**A**) Percent graft survival after Balbc÷B6 heterotopic heart transplant model in the setting of MR1 administration. Mice were injected intravenous with 0.25 mg tolerogen MR1 on day 0 then intraperitoneal on da ys 2 and 4 post-transplant. N=8 mice per group. (**B**) 5 weeks post transplant, H&E stained 10μm sections showed signs of CAV (vascular thickening) in *Axl+/+* recipients, however this was attenuated in *Axl-/-littermates*. To the right in a quantification of vascular thickening (N = 5 vs 5).

**Supplemental Figure 2.**
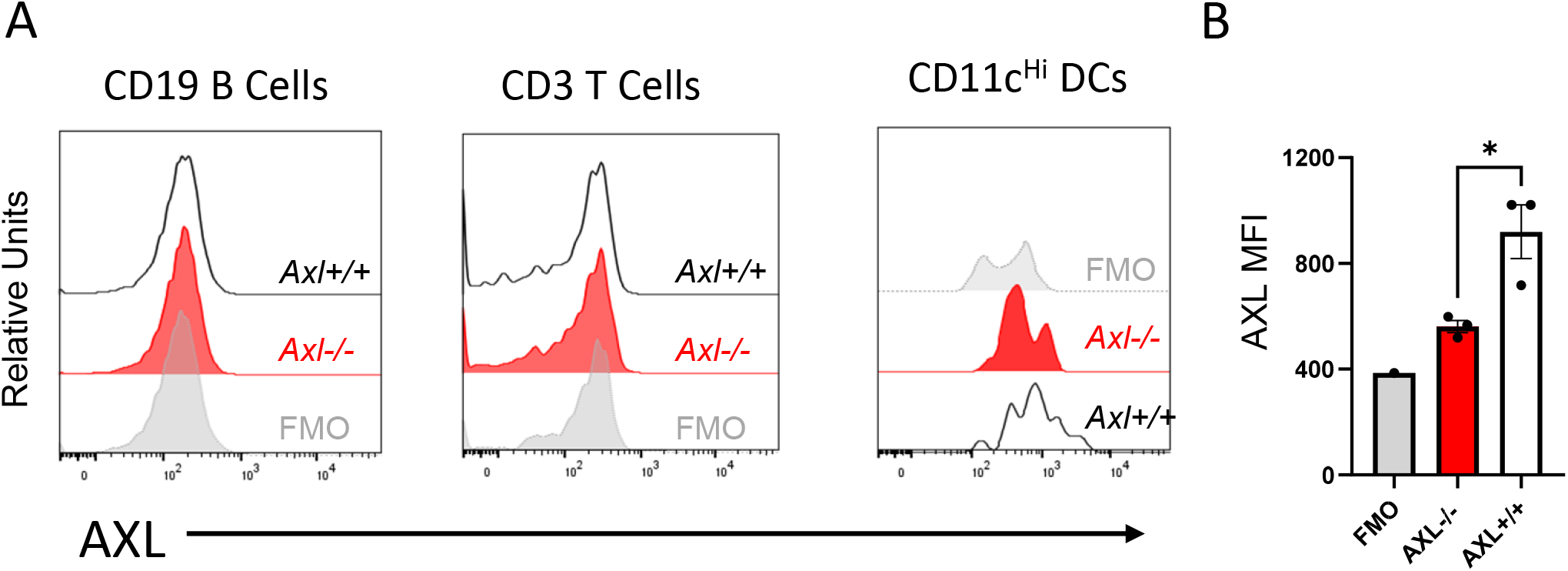
AXL protein on bone marrow-derived cells. B6 mice were exposed to a lethal dose of irradiation and consequently subjected to bone marrow cell transplantation by tail vein injection. After 4 weeks, reconstitution of immune cells was confirmed by flow cytometry. Subsets were further assessed for surface AXL staining. (**A**) CD45^+^CD19^+^ (B Cells) and CD45+CD3+ (T Cells) did not exhibit any appreciable AXL staining. Surface AXL was significantly detected in CD11c^Hi^CD11b^+^ (DCs) cells from recipients of *Axl+/+* bone marrow. (**B**) Quantification of AXL on DC-like cells in *Axl+/+* versus *Axl-/-* recipients. N=3 mice per group (* p<0.05).

**Supplemental Figure 3.**
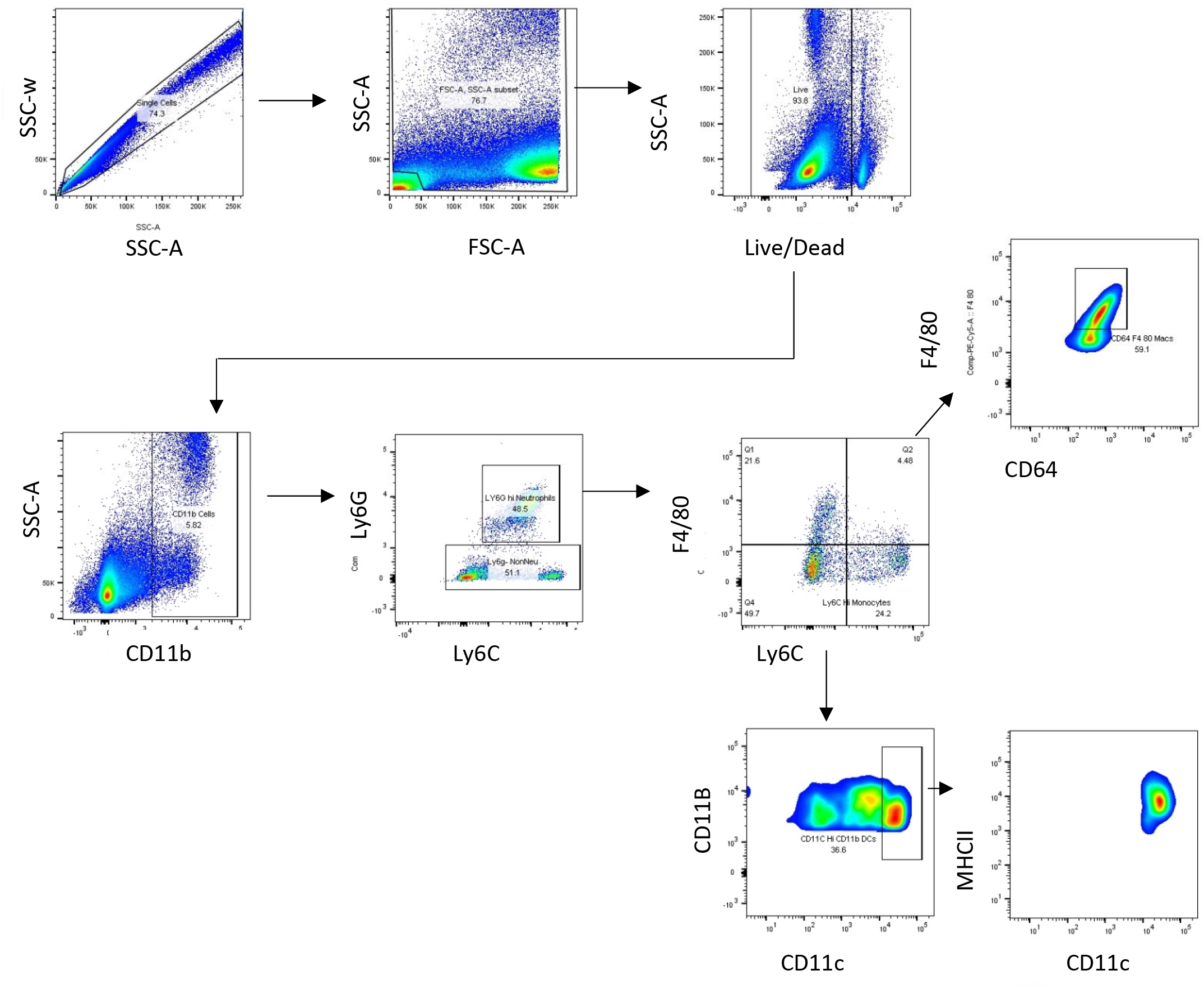
Gating strategy for myeloid subsets in recipient hearts and spleens. BM12 cardiac allografts and recipient spleens were extracted and subjected to flow cytometric analysis from B6 *Axl*+/+ versus *Axl*-/-recipients. After doublet exclusion and live/dead cell discrimination CD11b+ cells were selected. Neutrophils were then defined as CD11b^+^Ly6G^+^. Ly6G-were further distinguished as monocytes (CD11b^+^Ly6G^-^F4/80^-^Ly6C^lo^), macrophages (CD11b^+^Ly6G^-^F4/80^hi^CD64^+^Ly6C^lo^) and dendritic like cells (CD11b^+^Ly6G^-^F4/80^-^Ly6C^lo^F4/80^lo^CD11c^hi^).

**Supplemental Figure 4.**
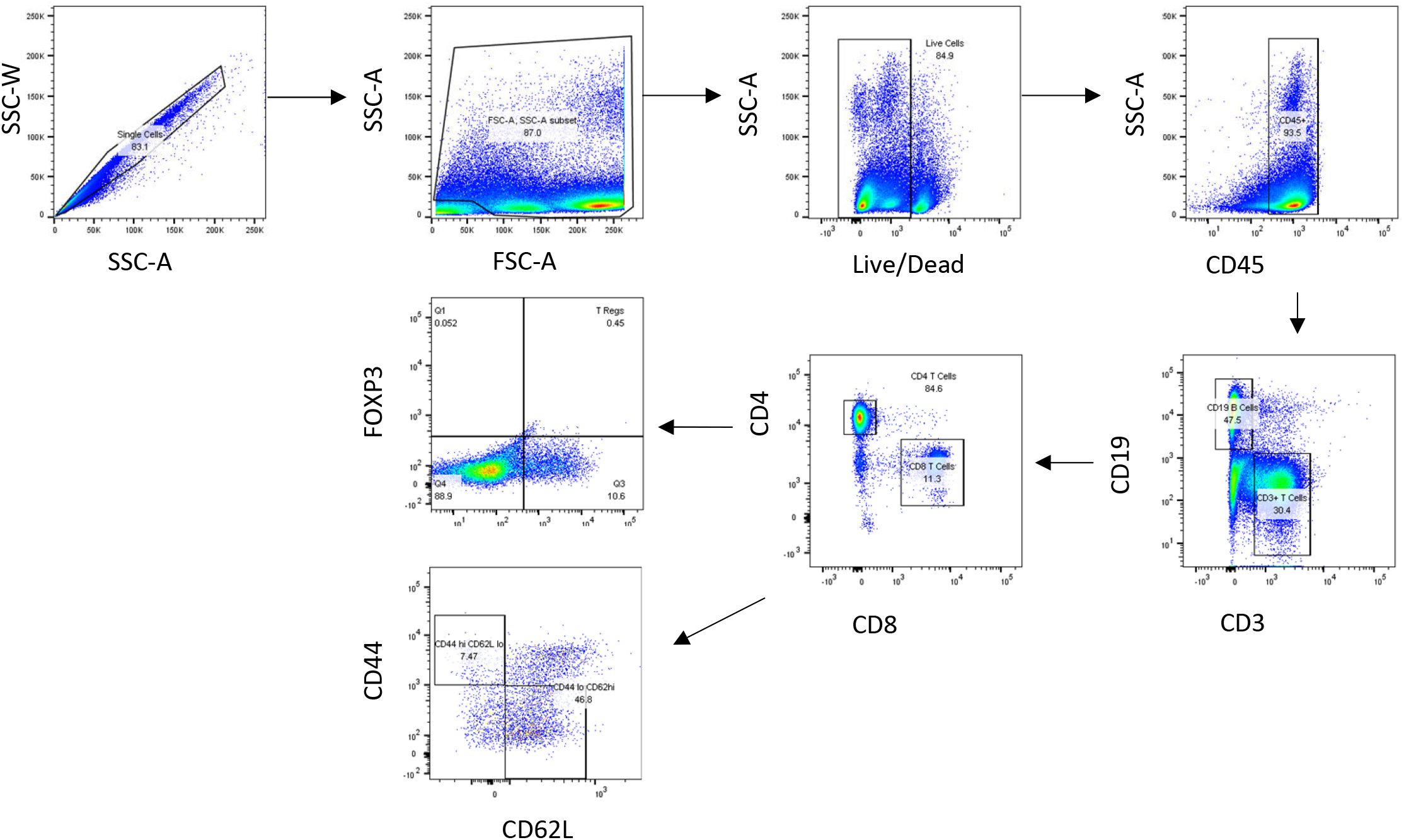
Gating strategy for lymphoid subsets in recipient hearts and spleens. BM12 cardiac allografts and recipient spleens were extracted and subjected to flow cytometric analysis from B6 *Axl*+/+ versus *Axl*-/-recipients. After doublet exclusion and live/dead cell discrimination CD45+ cells were selected. These were then separated by CD19 (B Cells) and CD3 (Tcell) staining. T cell subsets were further distinguished as CD4 T cells (CD45^+^CD3^+^CD8^-^CD4^+^) and CD8 T cells (CD45^+^CD3^+^CD4^-^CD8^+^). CD4 T cells were further identified as Tregs by possitive FoxP3^+^CD25^+^ staining, while memory T cells were identified as CD44^hi^ CD62L^lo^.

**Supplemental Figure 5.**
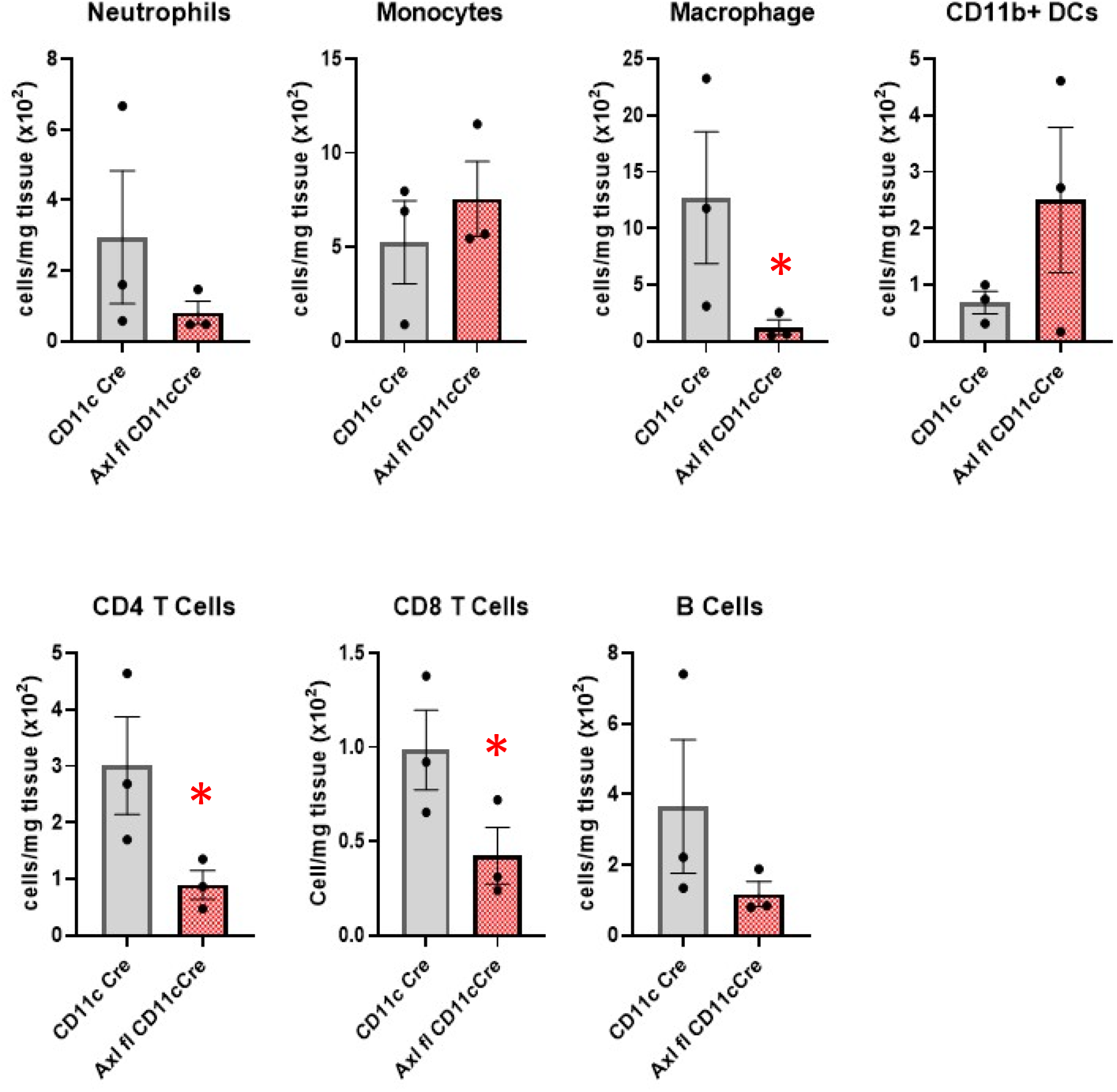
Reduced allograft inflammatory cell accumulation in the absence of *Axl* in CD11c+ cells. BM12 cardiac allografts were extracted and subjected to flow cytometric analysis from *B6;Axl^fl/fl^CD11c^cre^* versus *CD11c^cre^* recipients. Neutrophils were Ly6G^+^. Macrophages were F4/80^HI^. For T cell subsets, CD45^+^CD3^+^ cells were gated to distinguish CD4 and CD8 levels. N=3 mice per group. * P < 0.05

**Supplemental Figure 6.**
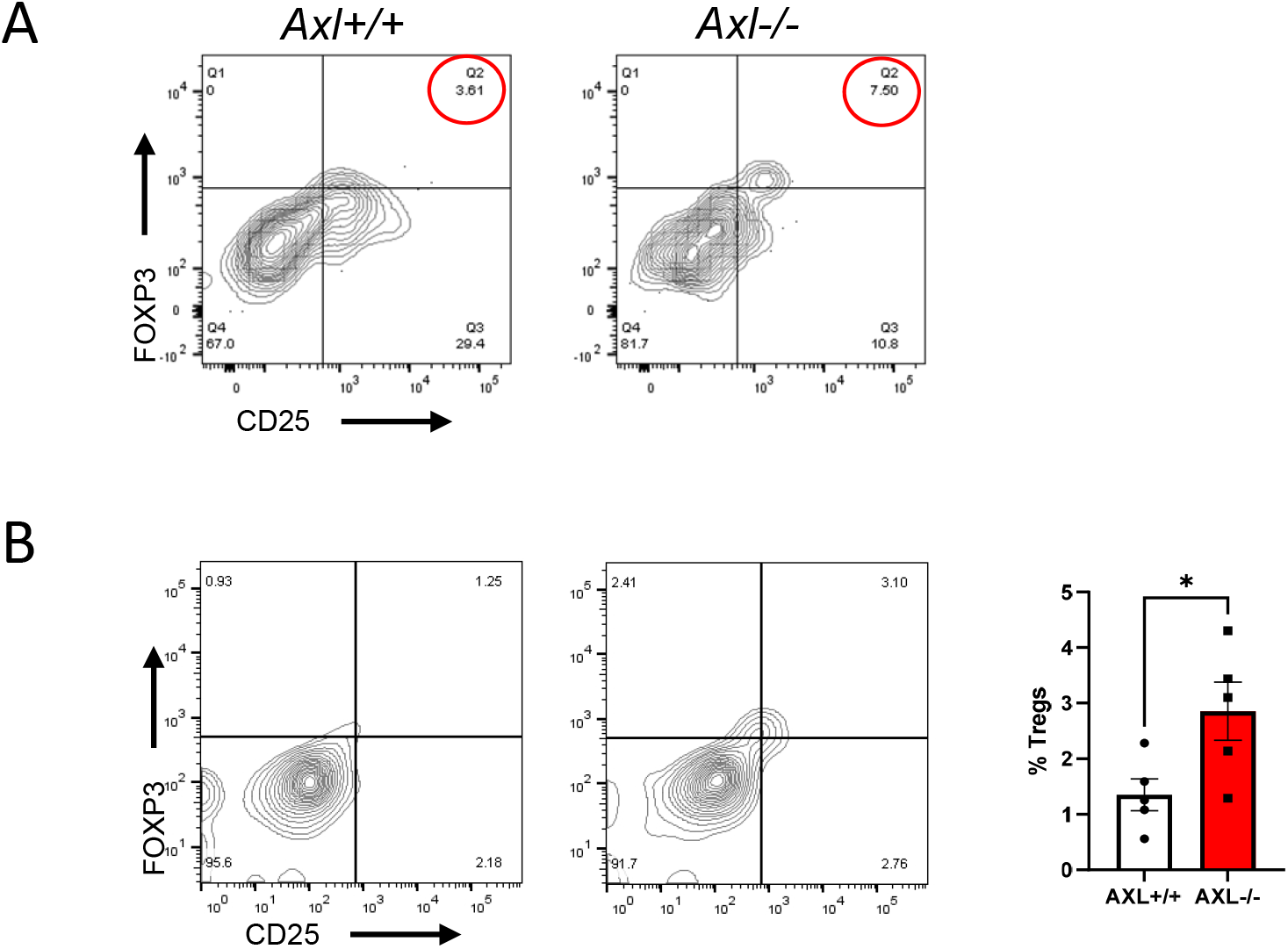
Evidence for elevated T regulatory cells post transplant in the absence of bone marrow-derived *Axl*. (**A**) *BM12* cardiac allografts were extracted and subjected to flow cytometric analysis from *B6 Axl+/+* versus *Axl-/-* recipients. T Cells were CD45^+^CD3^+^CD19^-^. T regs were further identified as CD4^+^CD25^+^Foxp3^+^. This is intra-graft analysis at 21days after transplant. (**B**) DC-like cells from *Axl+/+* versus *Axl-/-* bone marrow were assessed for their ability to promote Tregs in culture as indicated. N = 5 vs 5, p < 0.05.

**Supplemental Figure 7.**
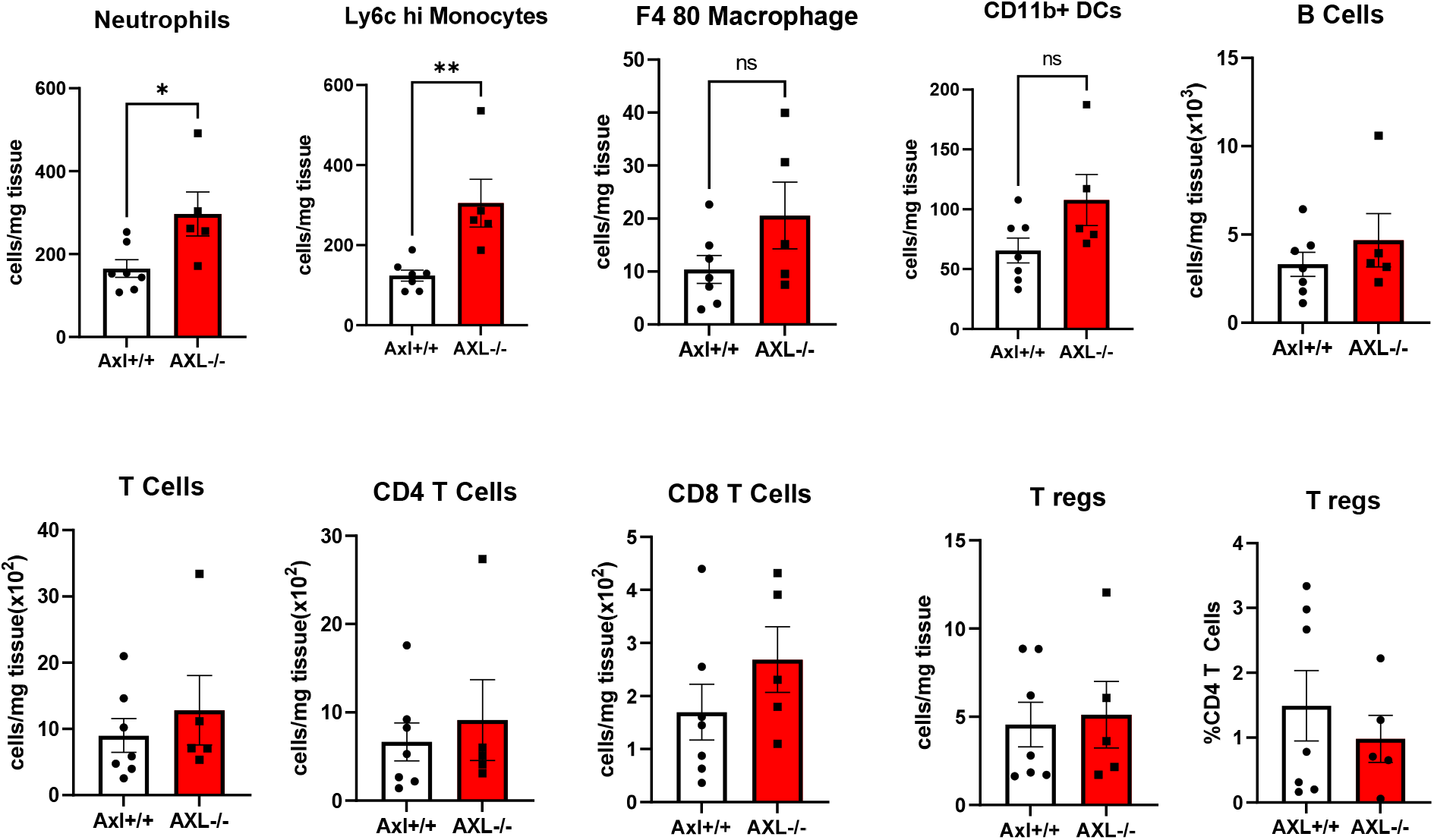
Flow cytometric analysis of recipient spleens. Analysis between *Axl+/+ and Axl-/-* groups as indicated. Ly6c^hi^ are monocytes and F4/80^+^ are macrophages and CD11b^+^ are DC-like cells. N= 8 vs 5 per group. * p < 0.05 ** p < 0.01.

**Supplemental Figure 8.**
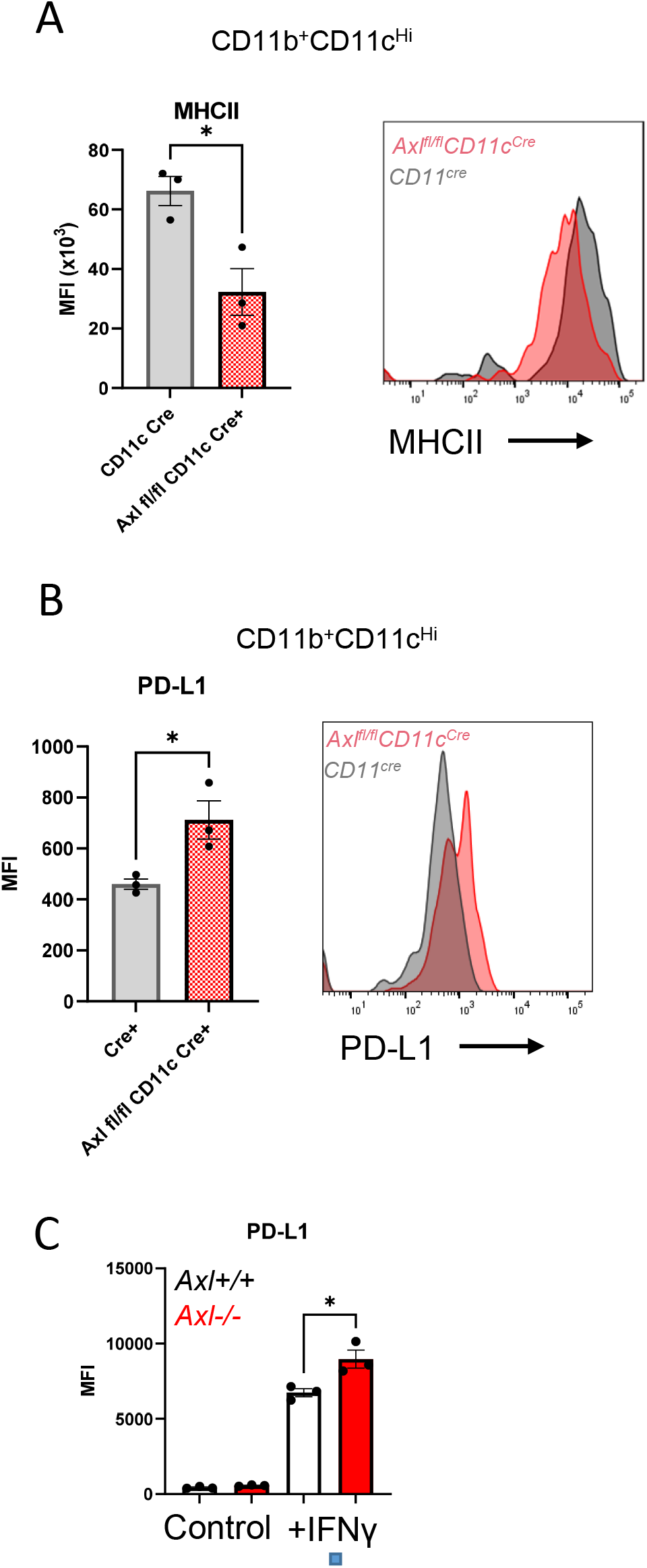
Evidence for elevated T regulatory cells post transplant in the absence of bone marrow-derived *Axl*. *BM12* cardiac allografts were extracted and subjected to flow cytometric analysis from *B6 Axl^fl/fl^CD11c^cre^* versus *CD11c^cre^* recipients. CD11+CD11c+ cells were assessed for (**A**) MHCII and (**B**) PD-L1 expression. (**C**) PD-L1 expression was assessed after IFNγ stimulation in bone marrow derived cells from *Axl+/+* and *Axl-/-* animals. N= 3 vs 3. P < 0.05.

**Supplemental Figure 9.**
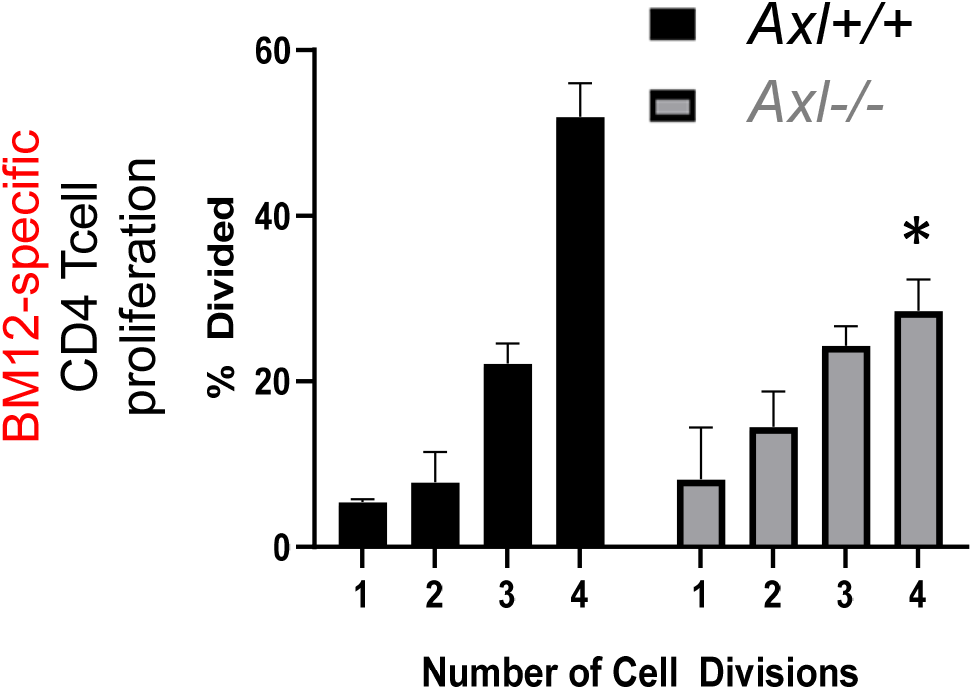
*Axl*-deficiency leads to depressed T cell activation. In culture, B6 dendritic cells were co-cultivated with irradiated BM12 cells and CFSE-labelled BM12-specific (ABM) T cells, as described in the Materials and Methods. CD4 Tcell proliferation was measured as a function of CFSE dilution. N=3 per variable. * = p < 0.05.

**Supplemental Figure 10.**
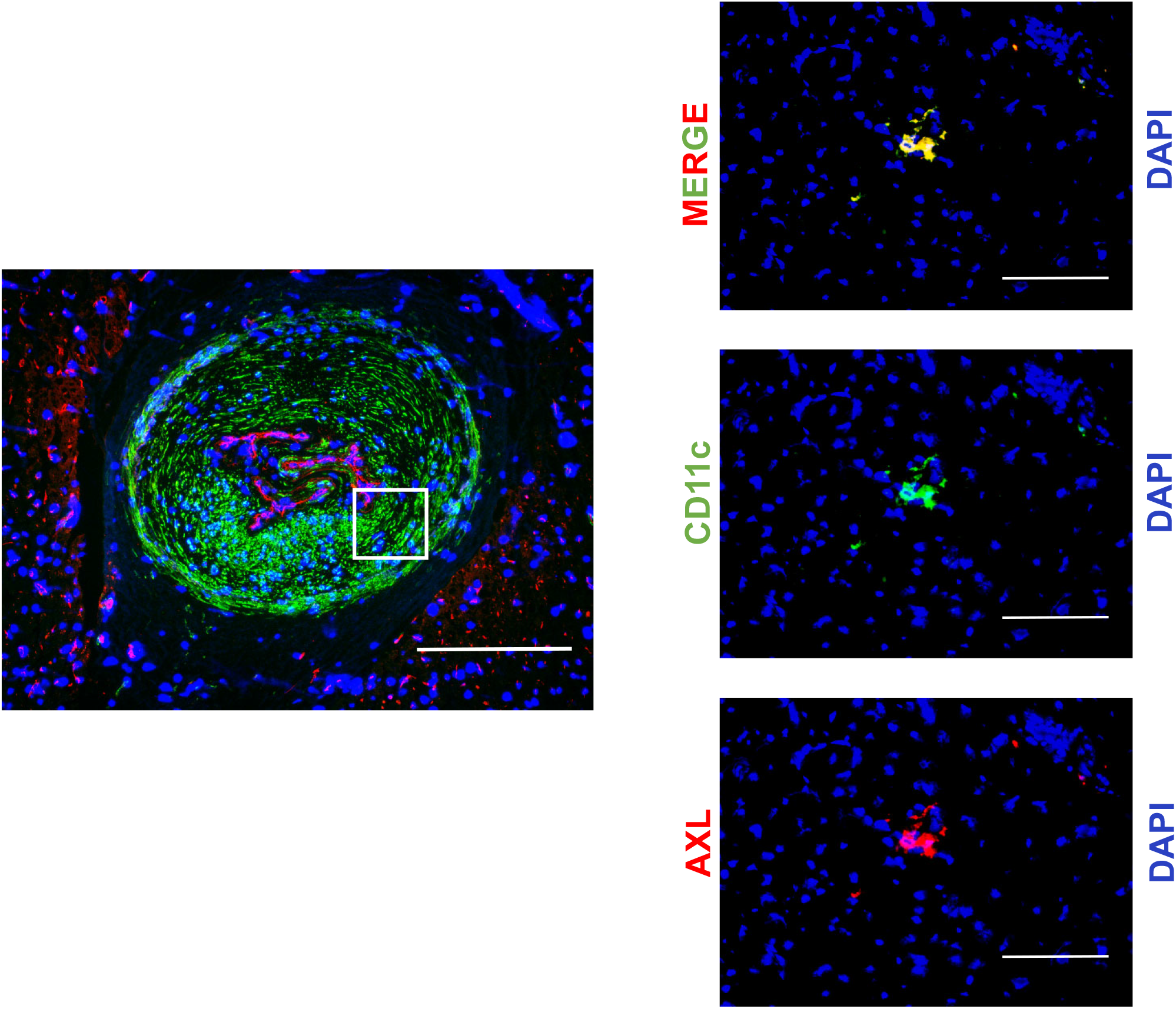
CD11c and alpha-smooth muscle actin allograft staining. Human cardiac sections were stained for CD11c high cells, alpha-smooth muscle actin (α-SMC) and DAPI for nuclei. Scale bars represent 400 and 100μm respectively.

**Supplemental Fig. 11.**
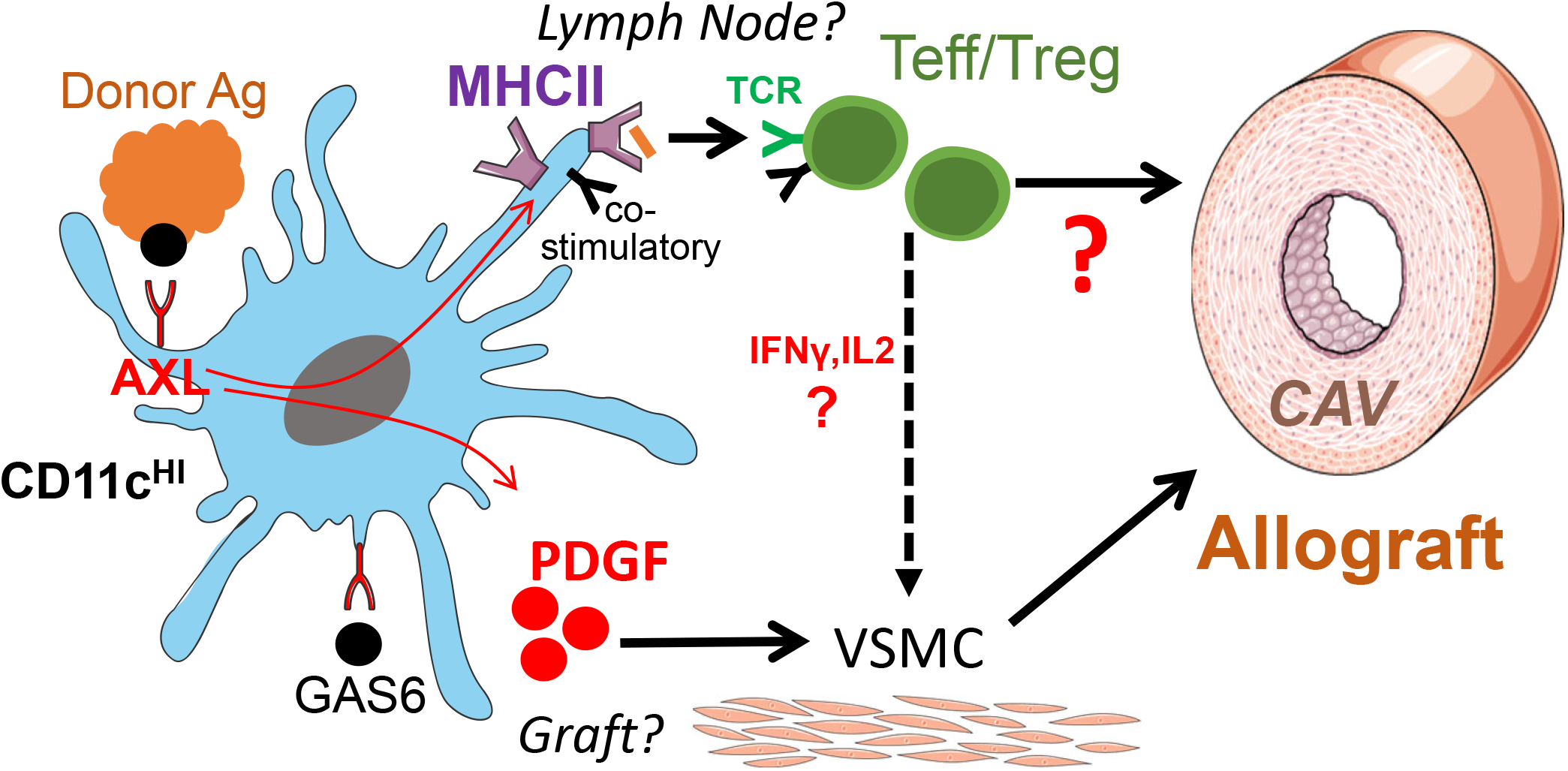
Working Model of Experimental Findings. Bone marrow AXL promotes vasculopathy and chronic rejection. In myeloid cells, AXL signaling regulates PDGF secretion to promote vascular smooth muscle cell (VSMC) proliferation and potentially contribute to cardiac allograft vasculopathy (CAV) from myeloid cells retained in the graft. AXL may also regulate antigen presentation and T cell activation via modulating MHCII and co-stimulatory molecule expression. This in turn could affects intra-graft T effector/T regulatory cell balance and further contribute to the inflammatory milieu. AXL may be activated by natural ligand GAS6 in the presence or absence of donor apoptotic cells.

## Notes

### Competing Interest Statement

The authors have declared no competing interest.

